# Evolutionary history of aquaporin-10 pseudogenization in Cetartiodactyla

**DOI:** 10.64898/2026.09.20.752922

**Authors:** Nodoka Nagai, Ayumi Nagashima, Akira Kato

**Affiliations:** School of Life Science and Technology, Institute of Science Tokyo, Yokohama, Japan

**Keywords:** Aquaporin-10, Cetartiodactyla, pseudogenization

## Abstract

Aquaporin-10 (AQP10) is an aquaglyceroporin that transports small, uncharged molecules, such as water and glycerol. In some species, such as mice and cattle, the AQP10 gene (*Aqp10*) is pseudogenized. In rodents, pseudogenization of *Aqp10* is limited to the Myomorpha suborder, while *Aqp10* remains intact in other rodent lineages. The order Cetartiodactyla encompasses diverse species, including Suina, Tylopoda, Ruminantia, and Whippomorpha. The distribution of intact *Aqp10* across the entire Cetartiodactyla and analyses estimating the timing of *Aqp10* pseudogenization have not been sufficiently investigated. In this study, we identified species within Cetartiodactyla that retain intact *Aqp10* and analyzed the detailed patterns of pseudogenization to elucidate the specifics of pseudogenization within this order. We collected *Aqp10* loci by BLAST and synteny analyses of publicly available genome databases of 51 cetartiodactyl species from 17 families. Dot plot and alignment analyses revealed that seudogenization of *Aqp10* occurred in 42 Cetartiodactyla species, including the Camelidae, warthog (Suidae), Antilocapridae, okapi (Giraffidae), Cervidae, Bovidae, Hippopotamidae, and Cetacea. In contrast, nine species retained an intact *Aqp10*: pig and babirusa (Suidae), peccary (Tayassuidae), mouse deeres (Tragulidae), giraffes (Giraffidae), and musk deers (Moschidae). The timing of pseudogenization was estimated by comparing patterns of exon deletions, frameshifts, and nonsense mutations in *Aqp10* pseudogenes. RT-PCR analysis of cattle and sheep tissues revealed that these species express *Aqp3* and *Aqp7* in their intestines, but not *Aqp10*. Collectively, these findings indicate that *Aqp10* pseudogenization occurred independently in multiple cetartiodactyl lineages and provide new insights into the evolutionary history of *Aqp10* loss within this order.

## 1. Introduction

Aquaglyceroporins are channels that transport small, uncharged molecules such as water and glycerol. There are four types of aquaglyceroporins in humans: AQP3, 7, 9, and 10. AQP10 is expressed in the human small intestine and adipocytes, contributing to nutrient absorption and glycerol supply to the blood (Hara-Chikuma and Verkman, 2006; Login and Nejsum, 2023; Rojek et al., 2008). An interesting feature of AQP10 is its high interspecies diversity. Early analyses revealed that the AQP10 gene (*Aqp10*) is pseudogenized in rodents, such as mice (Morinaga et al., 2002), and in artiodactyls, such as cattle (Tanaka et al., 2015). The absence of intact *Aqp10* in mice prevents the analysis of its physiological functions through observation of the knockout mouse phenotype. Detailed comparative genomic analysis of vertebrates by Yilmaz et al. revealed additional diversity in *Aqp10* (Yilmaz et al., 2020). The Actinopterygii (ray-finned fish) and the Chondrichthyes (cartilaginous fish) independently tandemly duplicated *Aqp10*, acquiring paralogs that are specific to their respective clades (Yilmaz et al., 2020). Although the specific physiological functions of these paralogs are unclear, biochemical analysis of their gene products shows significant differences in solute permeability. Ray-finned fish possess AQP10 paralogs with markedly different permeabilities to urea and boric acid (Imaizumi et al., 2024; Nagashima et al., 2025a; Nagashima et al., 2025c), whereas cartilaginous fish have AQP10s with distinct glycerol permeability (Hidaka et al., 2026; Nagashima and Kato, 2026).

Advances in next-generation sequencing technology in recent years have enabled the decoding and public release of numerous vertebrate genomes. Comparative genomic analysis using these public genome data has made it possible to observe the phenomenon of pseudogenization at higher resolution. For example, analysis of genomic data from 43 species across 13 rodent families revealed that *Aqp10* pseudogenization was observed in 13 species belonging to Myomorpha, while species belonging to Castorimorpha, Hystricomorpha, and Sciuromorpha retained intact *Aqp10* (Nagashima et al., 2025b). Pseudogenes are classified into unprocessed unitary, unprocessed duplicated, and processed types (Pink et al., 2011; Zhang et al., 2010). All *Aqp10* pseudogenes in rodents were unprocessed unitary pseudogenes (Nagashima et al., 2025b). These analyses have clarified the timing of *Aqp10* pseudogenization and the overall picture of *Aqp10* pseudogenization within rodents.

The order Cetartiodactyla comprises diverse organisms, including Suina (e.g., pigs), Tylopoda (camels, etc.), Ruminantia (e.g., cattle), and Whippomorpha (e.g., hippopotamuses and whales). This order encompasses 23 families and over 330 species (Zurano et al., 2019). Cetartiodactyla forms Laurasiatheria alongside Perissodactyla, Carnivora, Chiroptera, and others. Tanaka et al. analyzed the genome sequences of 14 Cetartiodactyla species and demonstrated that all 13 species except pigs have pseudogenized *Aqp10* (Tanaka et al., 2015). More recently, Rajput et al. analyzed the *Aqp* family in nine cetartiodactyl species and confirmed that the *Aqp10* had become pseudogenized in eight of these species, excluding pigs (Rajput et al., 2024). Currently, the National Center for Biotechnology Information (NCBI) dataset contains genomes from 190 distinct taxonomic IDs of artiodactyl species. In this study, we expanded the analysis to 51 cetartiodactyl species and systematically investigated the distribution of intact *Aqp10* and the evolutionary history of its pseudogenization. Our findings provide a comprehensive overview of *Aqp10* loss within Cetartiodactyla and offer new insights into the evolution of mammalian aquaglyceroporins.

## 2. Materials and Methods

### 2.1. Identification of *Aqp10* and pseudogenized *Aqp10* (*Aqp10p*) using Cetartiodactyla genome databases

The annotated *Aqp10* and *Aqp10p* genes and the unannotated genome regions for *Aqp10p* in Cetartiodactyla were collected by BLASTP or TBLASTN analysis using the National Center for Biotechnology Information (NCBI) BLAST server (https://blast.ncbi.nlm.nih.gov) (Johnson et al., 2008) and Ensembl genome browser (https://www.ensembl.org) (Martin et al., 2023) from the genome databases of the 51 Cetartiodactyla species listed in Table 1. The amino acid sequence of horse AQP10 (accession number, XP_001494035.1) was used as a query for the analyses.

**Table 1.**
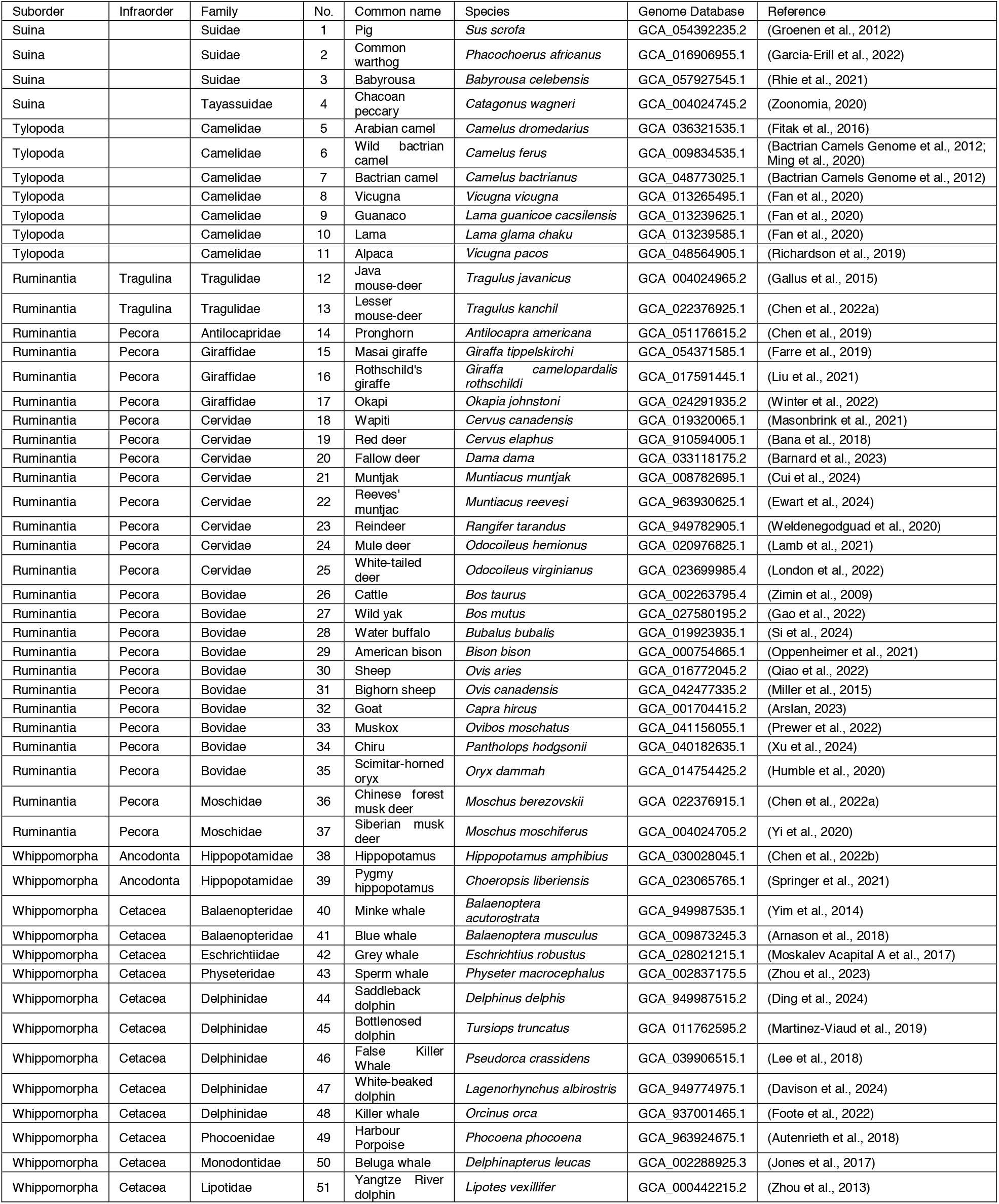
List of Cetartiodactyla species analyzed in this study.

| Suborder | Infraorder | Family | No. | Common name | Species | Genome Database | Reference |
| --- | --- | --- | --- | --- | --- | --- | --- |
| Suina |  | Suidae | 1 | Pig | *Sus scrofa* | GCA_054392235.2 | (Groenen et al., 2012) |
| Suina |  | Suidae | 2 | Common warthog | *Phacochoerus africanus* | GCA_016906955.1 | (Garcia-Erill et al., 2022) |
| Suina |  | Suidae | 3 | Babyrousa | *Babyrousa celebensis* | GCA_057927545.1 | (Rhie et al., 2021) |
| Suina |  | Tayassuidae | 4 | Chacoan peccary | *Catagonus wagneri* | GCA_004024745.2 | (Zoonomia, 2020) |
| Tylopoda |  | Camelidae | 5 | Arabian camel | *Camelus dromedarius* | GCA_036321535.1 | (Fitak et al., 2016) |
| Tylopoda |  | Camelidae | 6 | Wild bactrian camel | *Camelus ferus* | GCA_009834535.1 | (Bactrian Camels Genome et al., 2012; Ming et al., 2020) |
| Tylopoda |  | Camelidae | 7 | Bactrian camel | *Camelus bactrianus* | GCA_048773025.1 | (Bactrian Camels Genome et al., 2012) |
| Tylopoda |  | Camelidae | 8 | Vicugna | *Vicugna vicugna* | GCA_013265495.1 | (Fan et al., 2020) |
| Tylopoda |  | Camelidae | 9 | Guanaco | *Lama guanicoe cacsilensis* | GCA_013239625.1 | (Fan et al., 2020) |
| Tylopoda |  | Camelidae | 10 | Lama | *Lama glama chaku* | GCA_013239585.1 | (Fan et al., 2020) |
| Tylopoda |  | Camelidae | 11 | Alpaca | *Vicugna pacos* | GCA_048564905.1 | (Richardson et al., 2019) |
| Ruminantia | Tragulina | Tragulidae | 12 | Java mouse-deer | *Tragulus javanicus* | GCA_004024965.2 | (Gallus et al., 2015) |
| Ruminantia | Tragulina | Tragulidae | 13 | Lesser mouse-deer | *Tragulus kanchil* | GCA_022376925.1 | (Chen et al., 2022a) |
| Ruminantia | Pecora | Antilocapridae | 14 | Pronghorn | *Antilocapra americana* | GCA_051176615.2 | (Chen et al., 2019) |
| Ruminantia | Pecora | Giraffidae | 15 | Masai giraffe | *Giraffa tippelskirchi* | GCA_054371585.1 | (Farre et al., 2019) |
| Ruminantia | Pecora | Giraffidae | 16 | Rothschild's giraffe | *Giraffa camelopardalis rothschildi* | GCA_017591445.1 | (Liu et al., 2021) |
| Ruminantia | Pecora | Giraffidae | 17 | Okapi | *Okapia johnstoni* | GCA_024291935.2 | (Winter et al., 2022) |
| Ruminantia | Pecora | Cervidae | 18 | Wapiti | *Cervus canadensis* | GCA_019320065.1 | (Masonbrink et al., 2021) |
| Ruminantia | Pecora | Cervidae | 19 | Red deer | *Cervus elaphus* | GCA_910594005.1 | (Bana et al., 2018) |
| Ruminantia | Pecora | Cervidae | 20 | Fallow deer | *Dama dama* | GCA_033118175.2 | (Barnard et al., 2023) |
| Ruminantia | Pecora | Cervidae | 21 | Muntjak | *Muntiacus muntjak* | GCA_008782695.1 | (Cui et al., 2024) |
| Ruminantia | Pecora | Cervidae | 22 | Reeves' muntjak | *Muntiacus reevesi* | GCA_963930625.1 | (Ewart et al., 2024) |
| Ruminantia | Pecora | Cervidae | 23 | Reindeer | *Rangifer tarandus* | GCA_949782905.1 | (Weldenegodguad et al., 2020) |
| Ruminantia | Pecora | Cervidae | 24 | Mule deer | *Odocoileus hemionus* | GCA_020976825.1 | (Lamb et al., 2021) |
| Ruminantia | Pecora | Cervidae | 25 | White-tailed deer | *Odocoileus virginianus* | GCA_023699985.4 | (London et al., 2022) |
| Ruminantia | Pecora | Bovidae | 26 | Cattle | *Bos taurus* | GCA_002263795.4 | (Zimin et al., 2009) |
| Ruminantia | Pecora | Bovidae | 27 | Wild yak | *Bos mutus* | GCA_027580195.2 | (Gao et al., 2022) |
| Ruminantia | Pecora | Bovidae | 28 | Water buffalo | *Bubalus bubalis* | GCA_019923935.1 | (Si et al., 2024) |
| Ruminantia | Pecora | Bovidae | 29 | American bison | *Bison bison* | GCA_000754665.1 | (Oppenheimer et al., 2021) |
| Ruminantia | Pecora | Bovidae | 30 | Sheep | *Ovis aries* | GCA_016772045.2 | (Qiao et al., 2022) |
| Ruminantia | Pecora | Bovidae | 31 | Bighorn sheep | *Ovis canadensis* | GCA_042477335.2 | (Miller et al., 2015) |
| Ruminantia | Pecora | Bovidae | 32 | Goat | *Capra hircus* | GCA_001704415.2 | (Arslan, 2023) |
| Ruminantia | Pecora | Bovidae | 33 | Muskox | *Ovibos moschatus* | GCA_041156055.1 | (Prewer et al., 2022) |
| Ruminantia | Pecora | Bovidae | 34 | Chiru | *Pantholops hodgsonii* | GCA_040182635.1 | (Xu et al., 2024) |
| Ruminantia | Pecora | Bovidae | 35 | Scimitar-horned oryx | *Oryx dammah* | GCA_014754425.2 | (Humble et al., 2020) |
| Ruminantia | Pecora | Moschidae | 36 | Chinese forest musk deer | *Moschus berezovskii* | GCA_022376915.1 | (Chen et al., 2022a) |
| Ruminantia | Pecora | Moschidae | 37 | Siberian musk deer | *Moschus moschiferus* | GCA_004024705.2 | (Yi et al., 2020) |
| Whippomorpha | Ancodonta | Hippopotamidae | 38 | Hippopotamus | *Hippopotamus amphibius* | GCA_030028045.1 | (Chen et al., 2022b) |
| Whippomorpha | Ancodonta | Hippopotamidae | 39 | Pygmy hippopotamus | *Choeropsis liberiensis* | GCA_023065765.1 | (Springer et al., 2021) |
| Whippomorpha | Cetacea | Balaenopteridae | 40 | Minke whale | *Balaenoptera acutorostrata* | GCA_949987535.1 | (Yim et al., 2014) |
| Whippomorpha | Cetacea | Balaenopteridae | 41 | Blue whale | *Balaenoptera musculus* | GCA_009873245.3 | (Arnason et al., 2018) |
| Whippomorpha | Cetacea | Eschrichtiidae | 42 | Grey whale | *Eschrichtius robustus* | GCA_028021215.1 | (Moskalev Acapital A et al., 2017) |
| Whippomorpha | Cetacea | Physeteridae | 43 | Sperm whale | *Physeter macrocephalus* | GCA_002837175.5 | (Zhou et al., 2023) |
| Whippomorpha | Cetacea | Delphinidae | 44 | Saddleback dolphin | *Delphinus delphis* | GCA_949987515.2 | (Ding et al., 2024) |
| Whippomorpha | Cetacea | Delphinidae | 45 | Bottlenosed dolphin | *Tursiops truncatus* | GCA_011762595.2 | (Martinez-Viaud et al., 2019) |
| Whippomorpha | Cetacea | Delphinidae | 46 | False Killer Whale | *Pseudorca crassidens* | GCA_039906515.1 | (Lee et al., 2018) |
| Whippomorpha | Cetacea | Delphinidae | 47 | White-beaked dolphin | *Lagenorhynchus albirostris* | GCA_949774975.1 | (Davison et al., 2024) |
| Whippomorpha | Cetacea | Delphinidae | 48 | Killer whale | *Orcinus orca* | GCA_937001465.1 | (Foote et al., 2022) |
| Whippomorpha | Cetacea | Phocoenidae | 49 | Harbour Porpoise | *Phocoena phocoena* | GCA_963924675.1 | (Autenrieth et al., 2018) |
| Whippomorpha | Cetacea | Monodontidae | 50 | Beluga whale | *Delphinapterus leucas* | GCA_002288925.3 | (Jones et al., 2017) |
| Whippomorpha | Cetacea | Lipotidae | 51 | Yangtze River dolphin | *Lipotes vexillifer* | GCA_000442215.2 | (Zhou et al., 2013) |

### 2.2. Phylogenetic analysis of intact AQP10 in Cetartiodactyla

The nine amino acid sequences of intact AQP10 in Cetartiodactyla were aligned using ClustalW software (Chenna et al., 2003). Predicted AQP10 protein sequences for pig, Masai giraffe, and Chinese forest musk deer were retrieved from GenBank (accession numbers NP_001121926.1, XP_081350799.1, and XP_055259740.1, respectively). For the remaining five species, predicted protein sequences were manually reconstructed and are listed in Table S1. For comparison, AQP10 sequences from three perissodactyl species (horse, black rhinoceros, and Asiatic tapir), four carnivoran species (domestic cat, giant panda, Pacific walrus, and dog), human, and African savanna elephant were retrieved from GenBank and included in the analyses.

Their evolutionary histories were inferred using the maximum likelihood method and the Jones-Taylor-Thornton model (+Freq) (Jones et al., 1992). The tree with the highest log-likelihood (−2,340.72) is shown. The percentage of replicate trees in which the associated taxa clustered together (200 replicates) is shown next to the branches (Felsenstein, 1985). The initial tree for the heuristic search was selected by choosing the tree with the superior log-likelihood between a Neighbor-Joining (NJ) tree (Saitou and Nei, 1987) and a Maximum Parsimony (MP) tree. The NJ tree was generated using a matrix of pairwise distances computed using the Jones-Taylor-Thornton (1992) model (+Freq). We then selected the topology with the highest log-likelihood. The tree was drawn to scale with branch lengths measured as the number of substitutions per site. The analytical procedure encompassed 18 amino acid sequences with 302 positions in the final dataset. Evolutionary analyses were performed using the MEGA12 software (Kumar et al., 2024).

Synonymous and nonsynonymous nucleotide substitution rates (dS and dN, respectively) were calculated for *Aqp10* coding sequences. The coding sequences of *Aqp10* from six cetartiodactyl species, pig (*Sus scrofa*; NM_001128454.1), North Sulawesi babirusa (*Babyrousa celebensis*; Table S1), Chacoan peccary (*Catagonus wagneri*; Table S1), Java mouse-deer (*Tragulus javanicus*; Table S1), Masai giraffe (*Giraffa tippelskirchi*; XM_081494673.1), and Chinese forest musk deer (*Moschus berezovskii*; XM_055403765.1; Table S1) were aligned with those from three perissodactyl species, horse (*Equus caballus*; XM_001493985.5), black rhinoceros (*Diceros bicornis*; XM_058537482.1), and Asiatic tapir (*Tapirus indicus*; KAN7347935.1), and four carnivoran species, domestic cat (*Felis catus*; XM_019822500.3), giant panda (*Ailuropoda melanoleuca*; XM_002922533.4), Pacific walrus (*Odobenus rosmarus divergens*; XM_004403057.2), and dog (*Canis lupus familiaris*; XM_038543235.1). The nucleotide sequences of lesser mouse-deer (*Tragulus kanchil*), Rothschild’s giraffe (*Giraffa camelopardalis rothschildi*), and Siberian musk deer (*Moschus moschiferus*) were excluded from the analysis because they showed near-complete sequence identity to those of Java mouse-deer (*T. javanicus*), Masai giraffe (*G. tippelskirchi*), and Chinese forest musk deer (*M. berezovskii*) respectively. Gaps were manually adjusted to avoid frameshift artifacts, and the resulting alignment was used for the estimation of nucleotide substitution rates. The sequences were grouped according to the taxonomic orders of the corresponding species (Cetartiodactyla, Carnivora, and Perissodactyla). Mean within-group dS and dN values were calculated using the Nei-Gojobori (NG) method (Nei and Gojobori, 1986) with the complete-deletion option for gaps in MEGA12 (Kumar et al., 2024). The final alignment consisted of 13 coding nucleotide sequences and 299 codon positions.

### 2.3. Identification of nonsense mutation, frame shift, and exon deletion of *Aqp10p* in Cetartiodactyla

The protein coding regions of the exons in the intact *Aqp10* in Cetartiodactyla were aligned with the corresponding regions *Aqp10p* using ClustalW software, and the nonsense mutation and frame shift of *Aqp10p* were identified manually Fig. S1. Exon deletion was analyzed by dot plot analysis using the EMBOSS dotmatcher program (https://www.ebi.ac.uk/jdispatcher/emboss) (Madeira et al., 2024) with a window size of 20 and threshold score of 70 as previously described (Kato et al., 2024; Motoshima et al., 2023; Nagashima et al., 2025b) (Fig. S2). The genomic regions containing *Aqp10p* loci that were used for the dot plot analyses are listed in Table S2.

### 2.4. Synteny analysis of *Aqp10* or *Aqp10p* in Cetartiodactyla

Genome databases of the 51 Cetartiodactyla species listed in Table 1 were analyzed. Synteny analyses of genomic regions surrounding the *Aqp10p* locus (Table S3) were performed using the NCBI Genome Data Viewer (https://www.ncbi.nlm.nih.gov/genome/gdv/) (Rangwala et al., 2021) and Ensembl Genome Browser (https://www.ensembl.org) (Martin et al., 2023). The genome of the horse (*Equus caballus*; GCA_041296265.1) (Li et al., 2026), a representative species of the order Perissodactyla, was analyzed in the same manner as a comparator.

### 2.5. Semi-quantitative reverse transcription-polymerase chain reaction (RT-PCR)

Total RNAs of cattle, sheep, pig, and horse tissues were obtained from Zyagen (San Diego, CA, USA). First-strand complementary DNA was synthesized from 5 μg total RNA using the SuperScript IV First-Strand Synthesis System (Thermo Fisher Scientific) with Oligo(dT) primers and analyzed by RT-PCR as previously described (Tran et al., 2006). The cDNA was diluted eight times with nuclease-free water and used as the template for PCR with gene-specific primers (Table 2). Each reaction mixture (final volume, 12.5 μL) consisted of 0.25□μL cDNA (template), primers (individual final concentration, 0.25□μM), 6.25□μL GoTaq Green Master Mix (2×; Promega). The PCR conditions were as follows: initial denaturation at 94°C for 2 min, followed by 28 or 33□cycles at 94°C for 15□s (denaturation), 54°C for 30□s (annealing), 72°C for 1 min (extension), and a final extension at 72°C for 7 min. After amplification, the PCR mixture was diluted at 1:10 and was loaded 3 μL each onto a microchip electrophoresis system MCE-202 MultiNA (Shimadzu, Kyoto, Japan) using a DNA-12000 reagent kit (Shimadzu) according to the manufacturer’s instructions as previously described (Imaizumi et al., 2024; Motoshima et al., 2023; Nagashima et al., 2025b). Electrophoresis results were analyzed using the MultiNA Viewer software (Shimadzu).

**Table 2.**
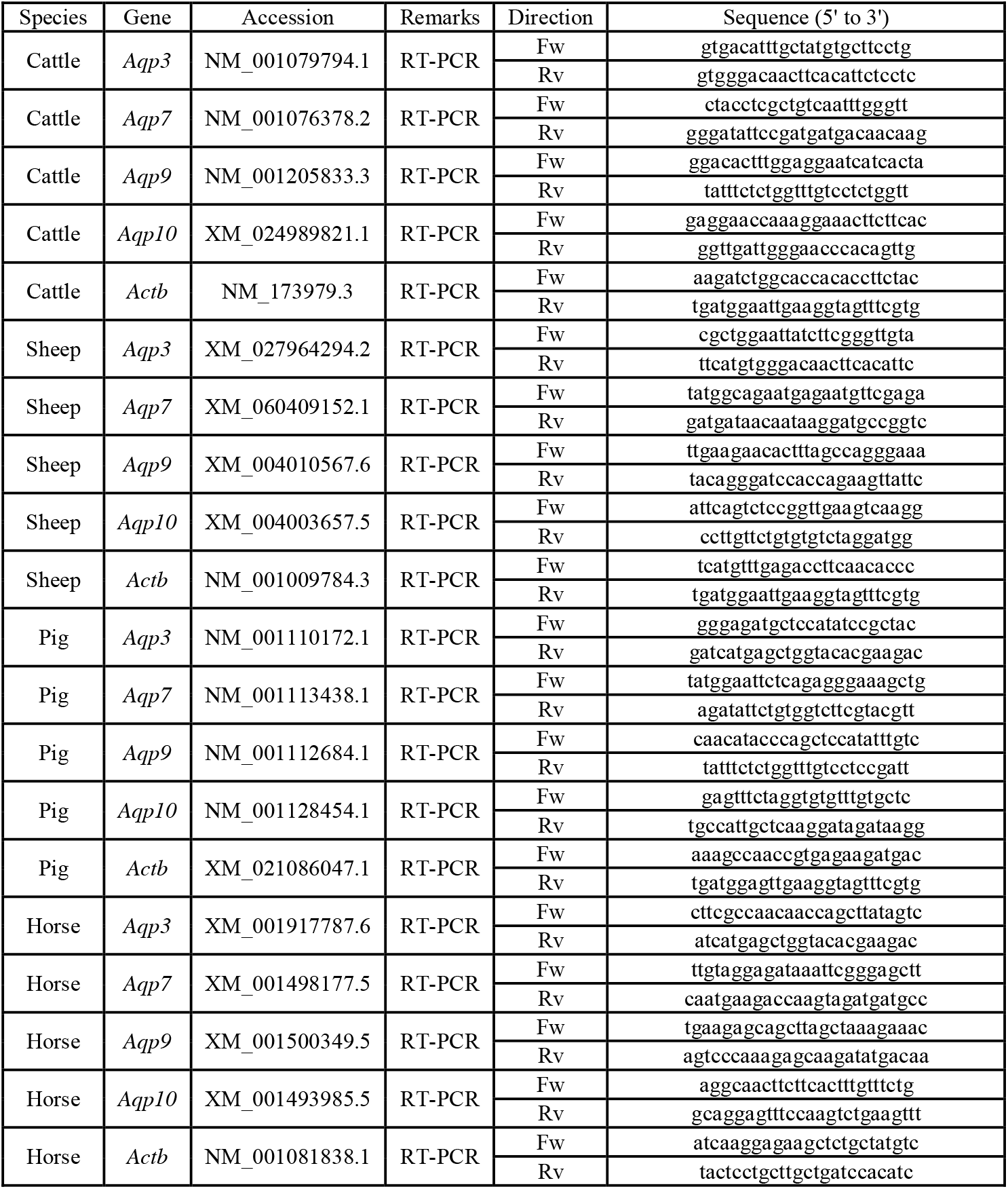
List of primers used for semiquantitative RT-PCR analysis.

## 3. Results

### 3.1. Retention of *Aqp10* in Cetartiodactyla

The phylogenetic relationships of the Cetartiodactyla species analyzed in this study are shown in Figure 1A. BLAST analysis of Cetartiodactyla identified intact AQP10 from nine species: Pig, North Sulawesi babirusa, Chacoan peccary, Java mouse-deer, Lesser mouse-deer, Masai giraffe, Rothschild’s giraffe, Chinese forest musk deer, and Siberian musk deer (Fig. 1A, species in blue). The predicted *Aqp10* coding sequence of the North Sulawesi babirusa encoded a protein that was 14 amino acid residues shorter at the C-terminus than that of pig AQP10. Because the C-terminal region of AQP10 shows low sequence conservation and substantial length variation among species, the babirusa *Aqp10* gene was classified as an intact gene. The predicted *Aqp10* sequences of Chinese forest musk deer and Siberian musk deer contained a putative translation initiation codon within an additional exon located upstream of exon 1, whereas no initiation codon corresponding to that found in exon 1 of other species was identified. Nevertheless, these genes were also classified as intact because an apparently intact open reading frame was retained.

**Fig. 1.**
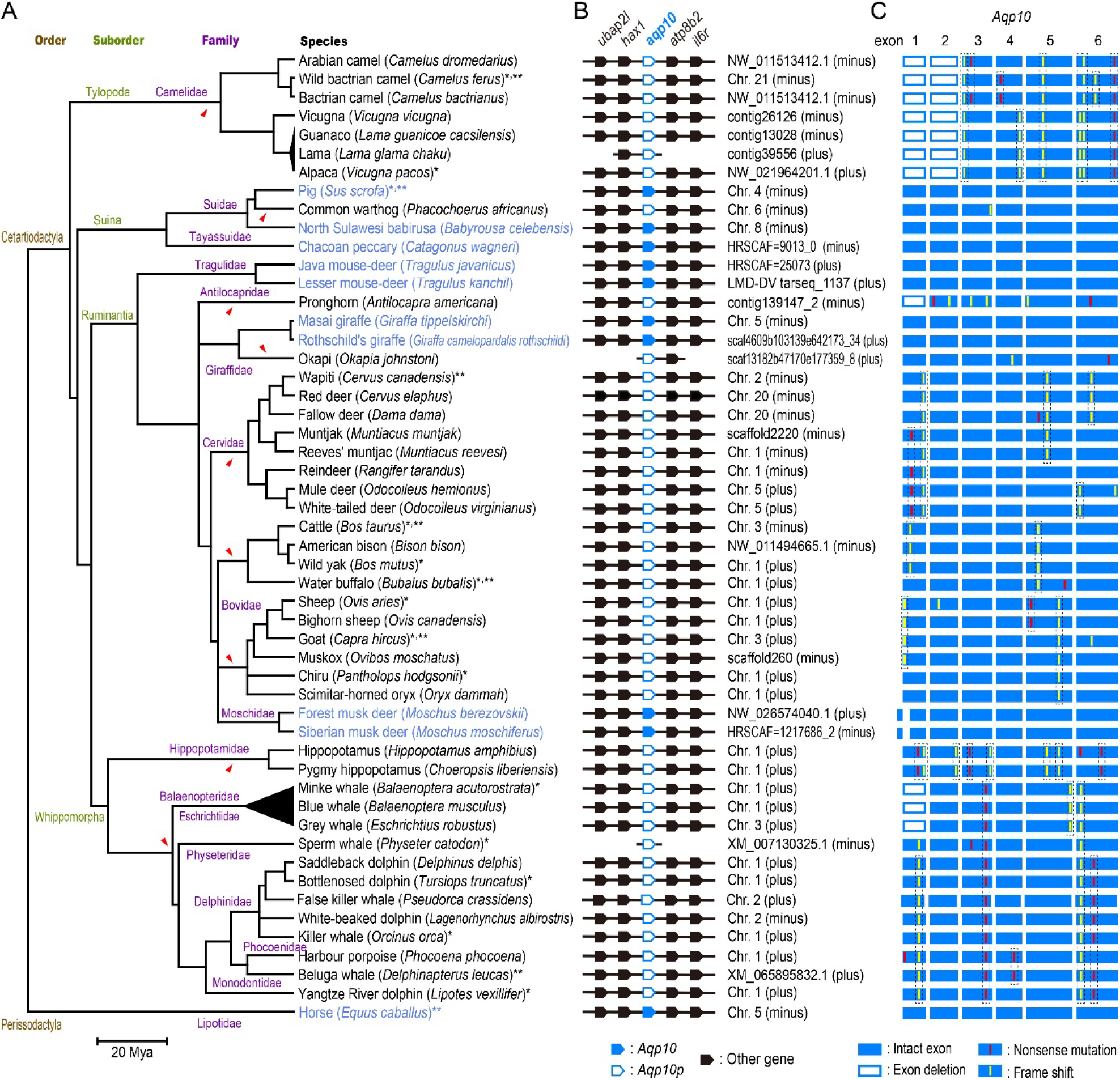
Pseudogenization of *Aqp10* in Cetartiodactyla. Presence of *Aqp10* or pseudogenized *Aqp10* (*Aqp10p*) in 51 cetartiodactyl species and horse are shown. (A) Phylogenetic relationship of 51 cetartiodactyl species analyzed in this study. Divergence times were retrieved from the TimeTree database (http://www.timetree.org/) (Kumar et al., 2022). Horse was analyzed as a related species not included in the Cetartiodactyla order. Asterisks (*) and double asterisks (**) indicate species that were also included in the studies of Tanaka et al. (Tanaka et al., 2015) and Rajput et al. (Rajput et al., 2024), respectively. Orange arrowheads indicate the inferred timing of pseudogenization. (B) Synteny of *Aqp10* and *Aqp10p* in various cetartiodactyl species. Synteny conservation of the region surrounding *Aqp10* and *Aqp10p* is shown. (C) *Aqp10* pseudogenization in cetartiodactyl species. Open reading frames divided into six exons are indicated by blue boxes. Deleted regions are indicated by open boxes. Nonsense and frameshift mutations are indicated by red and yellow bars, respectively. Supporting data for nucleotide sequence alignment and dot plot analyses are shown in Figs. S1 and S2.

In molecular phylogenetic analysis, the nine intact AQP10 identified from Cetartiodactyla formed a clade with AQP10 of other mammals (Fig. 2). Furthermore, the regions near the genomic locations encoding these proteins showed similarity in synteny with the horse genome region encoding AQP10 (Fig. 1B, filled blue boxes). These results confirm that the intact *Aqp10* genes identified from artiodactyls are the orthologs of *Aqp10*.

**Fig. 2.**
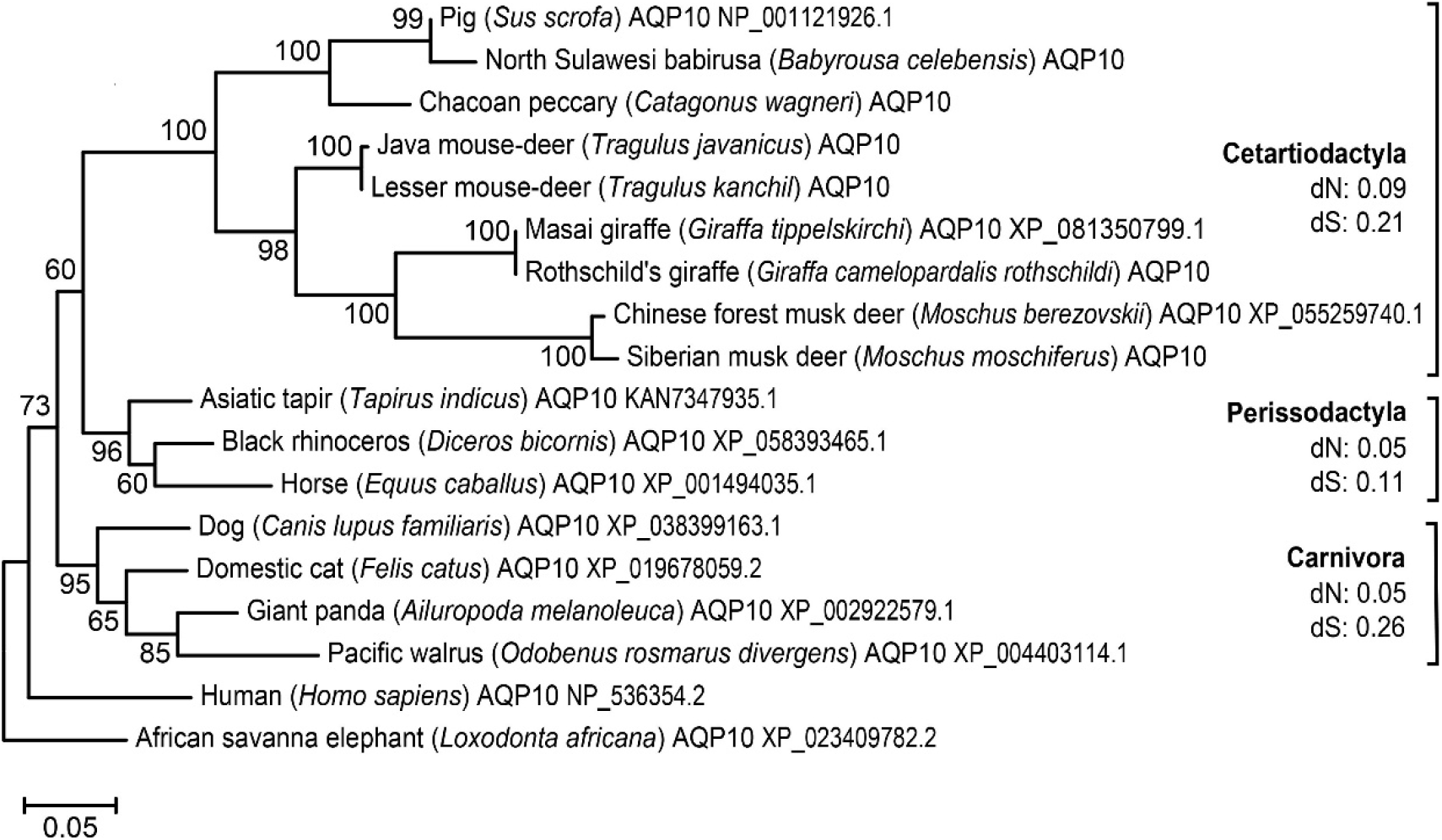
Phylogenetic analysis of intact AQP10 in Cetartiodactyla. A molecular phylogenetic tree was constructed from the amino acid sequences of intact *Aqp10* proteins identified in cetartiodactyl species and selected mammals using the maximum-likelihood (ML) method in MEGA12 (Kumar et al., 2024). Numbers at the nodes indicate bootstrap values. Mean dN and dS values for the *Aqp10* coding regions were calculated for Cetartiodactyla, Perissodactyla, and Carnivora and are shown to the right of the phylogenetic tree.

Synonymous and nonsynonymous substitution rates were analyzed for the *Aqp10* coding region, and the dN/dS ratio was calculated. The mean dN/dS ratio of *Aqp10* in cetartiodactyl species was 0.43. For comparison, the mean dN/dS ratios of *Aqp10* in three perissodactyl species and four carnivoran species were 0.45 and 0.19, respectively (Fig. 2). As all of these values were below 1, the intact *Aqp10* genes identified in cetartiodactyl species are inferred to have evolved under functional constraints, similar to *Aqp10* in Perissodactyla and Carnivora.

### 3.2. Pseudogenization of *Aqp10* in Cetartiodactyla

This study uses the term “pseudogene” in a broad sense, including all cases in which the gene is predicted to not encode a full-length water channel protein (Kato et al., 2024; Motoshima et al., 2023; Nagashima et al., 2025b; Tutar, 2012). BLAST analysis of 51 species within the Cetartiodactyla revealed that *Aqp10* in 42 species, excluding the aforementioned nine species, had become pseudogenized (Fig. 1). These include species previously reported by Tanaka et al. (Tanaka et al., 2015) and Rajput et al. (Rajput et al., 2024), indicated by asterisks and double asterisks, respectively, in Fig. 1A. Pseudogenization arises from mechanisms such as nonsense mutations, frameshifts, and exon deletions. Nonsense mutations and frame shifts were manually verified by aligning nucleic acid sequences corresponding to the *Aqp10p* region with intact Aqp10 exons (Fig. S1). Exon deletions were identified via dot plot analysis (Fig. S2). These results are schematically shown in Fig. 1C. Patterns of nonsense mutations, frame shifts, and exon deletions showed commonality within each lineage but differed between lineages. Within the lineages Camelidae, Cervidae, Bovinae, Caprinae, Hippopotamidae, Moschidae, and Cetaceans, unique, lineage-specific *Aqp10p* mutations were observed. Mutations in *Aqp10p* from the common warthog, pronghorn, and okapi showed no similarity to those in other lineages. The timing of predicted pseudogenization, as inferred from these results, is indicated by the orange arrowheads in Fig. 1A.

### 3.3. Expression of aquaglyceroporin genes in the intestines and kidneys of cattle, sheep, pig, and horse

In humans and guinea pigs, AQP10 is highly expressed in the small intestine (Ishibashi et al., 2002; Mobasheri et al., 2004; Nagashima et al., 2025b). Therefore, to identify aquaglyceroporins that might compensate AQP10 in the intestine, we confirmed the expression of bovine and ovine *Aqp-3, -7*, and *-9* using semiquantitative RT-PCR (Fig. 3). Simultaneously, we performed a similar analysis using primers specific for *Aqp10p*. In the bovine small intestine, *Aqp7* and *Aqp3* transcripts were detected in the duodenum, jejunum, and ileum. Similarly, *Aqp7* and *Aqp3* transcripts were detected in ovine duodenum, jejunum, and ileum. *Aqp9* transcripts were also detected in ovine jejunum. However, no amplification product was obtained with the *Aqp10p*-specific primers in either bovine or ovine tissues examined.

**Fig. 3.**
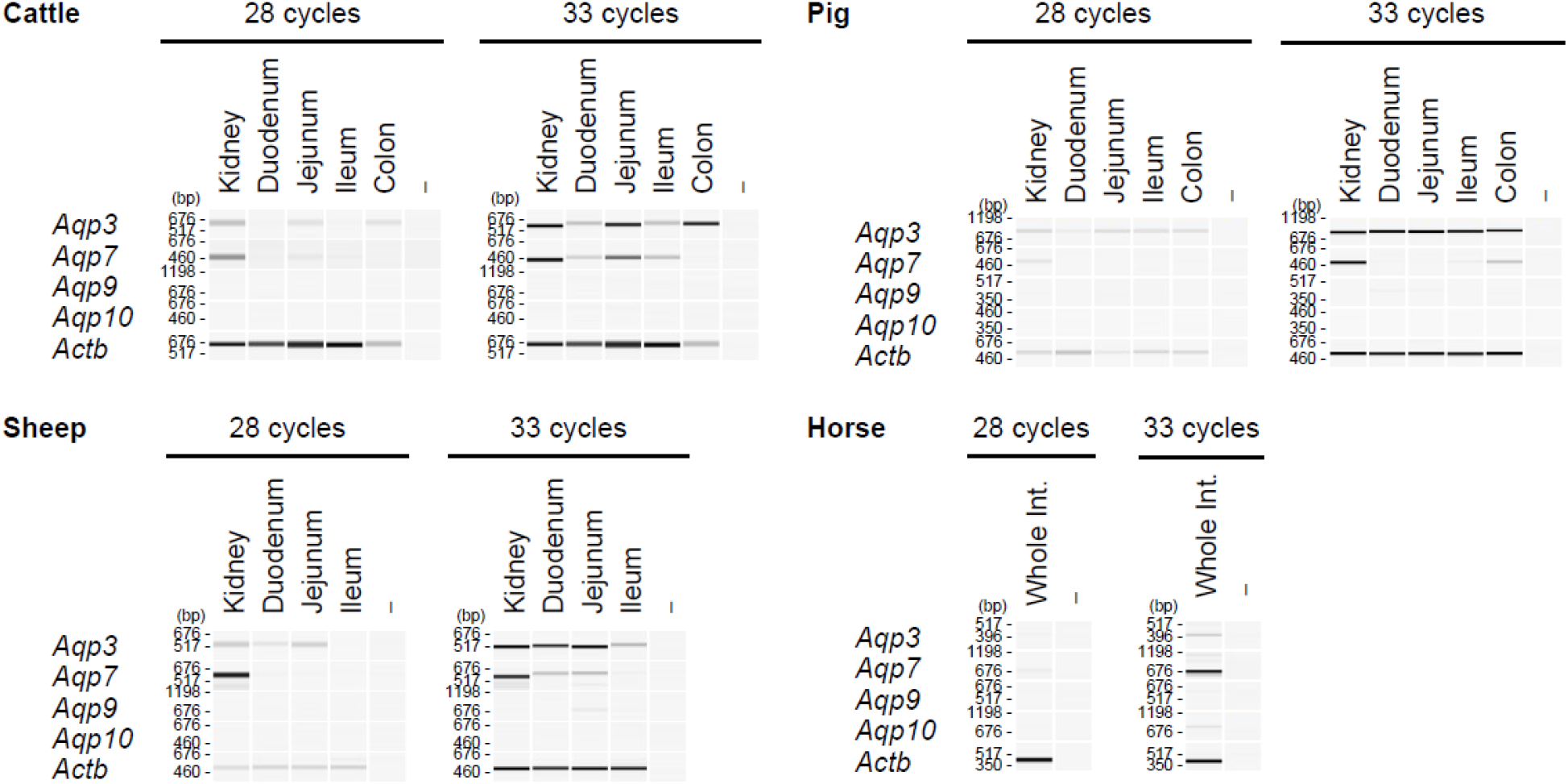
Intestinal and renal expression of aquaglyceroporins in cattle, sheep, pig, and horse. Expression profiles of *Aqp10* and related aquaglyceroporin genes in cattle, sheep, pig, and horse tissues were analyzed via semi-quantitative reverse transcription-polymerase chain reaction (RT-PCR). Pseudo-gel images of PCR products were generated using a microchip electrophoresis system. β-actin (Actb) was used as an internal control. Whole images of the gels are shown in Fig. S3.

For comparison, intestinal expression of aquaglyceroporin genes was also examined in Aqp10-retaining Cetartiodactyla (pigs) and Perissodactyla (horses) (Fig. 3). In pig, *Aqp3* transcripts were detected in the duodenum, jejunum, and ileum. *Aqp7* transcripts were detected in the ileum, whereas *Aqp9* transcripts were detected in the duodenum and jejunum. In contrast, Aqp10 expression was not detected in the pig tissues examined. In horse, RNA prepared from the whole intestine showed detectable expression of *Aqp-3, -7*, and *-10* was observed.

## 4. Discussion

Previous reports on Cetartiodactyla species (Rajput et al., 2024; Tanaka et al., 2015) indicated that *Aqp10* in cetartiodactyls other than pigs had become pseudogenized. In this study, we expanded the analysis to 51 species and confirmed this trend while making several new discoveries. Within Suina, the pig and North Sulawesi babirusa in Suidae and Chacoan peccary in Tayassuidae retained intact *Aqp10*. However, the common warthog’s *Aqp10* contained a frameshift mutation in exon 2, suggesting it is pseudogenized. Thus, although *Aqp10* is generally intact in Suina, exceptions exist.

All analyzed species belonging to Camelidae and Whippomorpha possessed pseudogenized *Aqp10*. In Camelidae, several disruptive mutations in *Aqp10p*, including deletions of exons 1 and 2 and frameshift-inducing insertions in exons 3, 5, and 6, were shared among all camelid species examined. This pattern suggests that *Aqp10* pseudogenization likely occurred in the common ancestor of Camelidae. In contrast, within the Whippomorpha lineage, multiple shared nonsense mutations and frameshift mutations were identified within Hippopotamidae and Cetacea, respectively, whereas no disruptive mutations were shared between the two clades. These findings suggest that pseudogenization of *Aqp10* occurred independently in the common ancestors of Hippopotamidae and Cetacea after their divergence.

Although most ruminant species possessed pseudogenized *Aqp10*, intact copies were retained in two mouse-deer species, two giraffe species, and two musk deer species, indicating that a small number of ruminant lineages have maintained a functional copy. A comparative analysis of exon deletions, nonsense mutations, and frameshift mutations in *Aqp10* pseudogenes revealed that several mutations were shared within individual ruminant clades, suggesting that these mutations were acquired in their respective common ancestors. Taken together, these observations suggest that *Aqp10* pseudogenization arose independently in multiple cetartiodactyl lineages (Fig. 1).

The *Aqp10* pseudogenes identified in Cetartiodactyla were located at loci syntenic to those harboring intact *Aqp10* in other species and retained remnants of the ancestral exon-intron structure. In addition, no *Aqp10* paralogs were identified elsewhere in the genome. Together, these characteristics indicate that cetartiodactyl *Aqp10p* genes are best classified as unprocessed unitary pseudogenes rather than unprocessed duplicated or processed pseudogenes (Pink et al., 2011; Zhang et al., 2010).

In species that retain intact *Aqp10*, the intestine is generally considered a major site of expression. The distribution of intact and pseudogenized *Aqp10* among cetartiodactyl lineages raises the possibility that digestive physiology associated feeding adaptations may have influenced the retention or loss of *Aqp10* during cetartiodactyl evolution. However, the limited number of species retaining intact *Aqp10* and the presence of such species within Ruminantia suggest that digestive anatomy alone cannot fully explain the observed pattern. According to the classification of Zurano et al., Suina diverged first within Cetartiodactyla, while Tylopoda, Ruminantia, and Whippomorpha form a single clade. Suina possess a single stomach, whereas Tylopoda, Ruminantia, and Whippomorpha possess multi-chambered stomachs. Ruminants, in particular, possess a multi-chambered stomach. They digest dietary fiber, such as cellulose, through rumination and absorb it as short-chain fatty acids, like propionic acid (Baldwin VI et al., 2004; Russell et al., 2009). Ruminants have also evolved the ability to secrete urea into their saliva and digestive tracts via urea transporters. This enables them to absorb amino acids produced by microorganisms as nutrients (Baldwin VI et al., 2004; Stewart and Smith, 2005). Given that several species belonging to the monogastric Suina possess intact *Aqp10*, and that *Aqp10*—which functions as a glycerol-permeable aquaglyceroporin—is pseudogenized in most ruminants, these patterns are consistent with the possibility that the selective pressures maintaining *Aqp10* function were relaxed in many ruminant lineages.

In addition to AQP10, mammals possess AQP-3, -7, and -9, which exhibit similar channel activities. In absorptive epithelial cells, AQP7 and AQP10 are predominantly localized to the apical membrane, whereas AQP3 is localized to the basolateral membrane (Ishibashi et al., 2000; Mobasheri et al., 2004; Sohara et al., 2006). In rodents lacking *Aqp10, Aqp7* expression has been observed in the intestines, suggesting that *Aqp7* may substitute for the function of *Aqp10* (Nagashima et al., 2025b). The present RT-PCR analysis detected *Aqp7* expression in the intestines of cattle and sheep (Fig. 2). These results suggest that *Aqp7* may substitute for *Aqp10* function in the intestines of ruminants lacking *Aqp10*. Interestingly, pigs retain an intact *Aqp10* coding sequence, yet intestinal expression was not detected. Together with similar observations in rats (Koyama et al., 1999; Laforenza, 2012; Nagashima et al., 2025b), this finding suggests that maintenance of an intact *Aqp10* sequence does not necessarily imply active intestinal expression. In these species, *Aqp7* may fulfill physiological roles that are otherwise served by *Aqp10* in other mammals.

## CRediT authorship contribution statement

**Nodoka Nagai:** Conceptualization; Formal analysis; Investigation; Data Curation; Writing - Original Draft; Visualization; Project administration. **Ayumi Nagashima:** Investigation; Resources; Funding acquisition. **Akira Kato:** Conceptualization; Methodology; Validation; Formal analysis; Investigation; Resources; Data Curation; Writing - Original Draft; Writing - Review & Editing; Visualization; Supervision; Project administration; Funding acquisition.

## Acknowledgments

We thank the Open Research Facilities for Life Science and Technology at the Institute of Science, Tokyo, for their technical assistance.

## Data availability

Data will be made available upon reasonable request.

## References

Arnason, U., Lammers, F., Kumar, V., Nilsson, M.A., Janke, A., 2018. Whole-genome sequencing of the blue whale and other rorquals finds signatures for introgressive gene flow. Sci Adv 4, eaap9873.

Arslan, M., 2023. Whole-genome sequencing and genomic analysis of Norduz goat (Capra hircus). Mamm. Genome 34, 437–448.

Autenrieth, M., Hartmann, S., Lah, L., Roos, A., Dennis, A.B., Tiedemann, R., 2018. High-quality whole-genome sequence of an abundant Holarctic odontocete, the harbour porpoise (Phocoena phocoena). Mol Ecol Resour 18, 1469–1481.

Bactrian Camels Genome, S., Analysis, C., Jirimutu, Wang, Z., Ding, G., Chen, G., Sun, Y., Sun, Z., Zhang, H., Wang, L., Hasi, S., Zhang, Y., Li, J., Shi, Y., Xu, Z., He, C., Yu, S., Li, S., Zhang, W., Batmunkh, M., Ts, B., Narenbatu, Unierhu, Bat-Ireedui, S., Gao, H., Baysgalan, B., Li, Q., Jia, Z., Turigenbayila, Subudenggerile, Narenmanduhu, Wang, Z., Wang, J., Pan, L., Chen, Y., Ganerdene, Y., Dabxilt, Erdemt, Altansha, Altansukh, Liu, T., Cao, M., Aruuntsever, Bayart, Hosblig, He F., Zha-ti, A., Zheng, G., Qiu, F., Sun, Z., Zhao, L., Zhao, W., Liu, B., Li, C., Chen, Y., Tang, X., Guo, C., Liu, W., Ming, L., Temuulen Cui, A., Li, Y., Gao, J., Li, J., Wurentaodi Niu, S., Sun, T., Zhai, Z., Zhang, M., Chen, C., Baldan, T., Bayaer, T., Li, Y., Meng, H., 2012. Genome sequences of wild and domestic bactrian camels. Nat Commun 3, 1202.

Baldwin VI, R.L., McLeod, K.R., Klotz, J.L., Heitmann, R.N., 2004. Rumen development, intestinal growth and hepatic metabolism in the pre-and postweaning ruminant. J. Dairy Sci. 87, E55–E65.

Bana, N.A., Nyiri, A., Nagy, J., Frank, K., Nagy, T., Steger, V., Schiller, M., Lakatos, P., Sugar, L., Horn, P., Barta, E., Orosz, L., 2018. The red deer Cervus elaphus genome CerEla1.0: sequencing, annotating, genes, and chromosomes. Mol Genet Genomics 293, 665–684.

Barnard, R.K., Smith, J.A., Yuan, N., Liu, F., Hadi, S.S., 2023. An announcement of a new genome sequence available for Dama dama (fallow deer). Forensic Sci Int Anim Environ 4, 100074.

Chen, C., Yin, Y., Li, H., Zhou, B., Zhou, J., Zhou, X., Li, Z., Liu, G., Pan, X., Zhang, R., Lin, Z., Chen, L., Qiu, Q., Zhang, Y.E., Wang, W., 2022a. Ruminant-specific genes identified using high-quality genome data and their roles in rumen evolution. Sci Bull (Beijing) 67, 825–835.

Chen, L., Li, Z., Wu, B., Zhou, B., Heller, R., Zhou, J., Wang, K., Lin, Z., Wu, D., Qiu, Q., 2022b. Progressive evolution of secondary aquatic adaptation in hippos and cetaceans. Cell Discov 8, 134.

Chen, L., Qiu, Q., Jiang, Y., Wang, K., Lin, Z., Li, Z., Bibi, F., Yang, Y., Wang, J., Nie, W., Su, W., Liu, G., Li, Q., Fu, W., Pan, X., Liu, C., Yang, J., Zhang, C., Yin, Y., Wang, Y., Zhao, Y., Zhang, C., Wang, Z., Qin, Y., Liu, W., Wang, B., Ren, Y., Zhang, R., Zeng, Y., da Fonseca, R.R., Wei, B., Li, R., Wan, W., Zhao, R., Zhu, W., Wang, Y., Duan, S., Gao, Y., Zhang, Y.E., Chen, C., Hvilsom, C., Epps, C.W., Chemnick, L.G., Dong, Y., Mirarab, S., Siegismund, H.R., Ryder, O.A., Gilbert, M.T.P., Lewin, H.A., Zhang, G., Heller, R., Wang, W., 2019. Large-scale ruminant genome sequencing provides insights into their evolution and distinct traits. Science 364.

Chenna, R., Sugawara, H., Koike, T., Lopez, R., Gibson, T.J., Higgins, D.G., Thompson, J.D., 2003. Multiple sequence alignment with the Clustal series of programs. Nucleic Acids Res 31, 3497–3500.

Cui, Y., Lv, Y., Li, J., Cai, M., Liu, X., Xu, Z., Liu, H., 2024. A haplotype-resolved and chromosome-scale genome assembly of Hainan muntjac (Muntiacus nigripes). Sci Data 11, 1395.

Davison, N.J., Morin, P., Wellcome Sanger Institute Tree of Life Management, S., Laboratory, t., Wellcome Sanger Institute Scientific Operations: Sequencing, O., Wellcome Sanger Institute Tree of Life Core Informatics, t., Tree of Life Core Informatics, c., Darwin Tree of Life, C., 2024. The genome sequence of the white-beaked dolphin, Lagenorhynchus albirostris (Gray, 1846). Wellcome Open Res 9, 687.

Ding, K., Xu, Q., Zhao, L., Li, Y., Li, Z., Shi, W., Zeng, Q., Wang, X., Zhang, X., 2024. Chromosome-level genome provides insights into environmental adaptability and innate immunity in the common dolphin (Delphinus delphis). BMC Genomics 25, 373.

Ewart, N., Wawman, D.C., University of, O., Wytham Woods Genome Acquisition, L., Darwin Tree of Life Barcoding, c., Wellcome Sanger Institute Tree of Life Management, S., Laboratory, t., Wellcome Sanger Institute Scientific Operations: Sequencing, O., Wellcome Sanger Institute Tree of Life Core Informatics, t., Tree of Life Core Informatics, c., Darwin Tree of Life, C., 2024. The genome sequence of Reeves’ muntjac Muntiacus reevesi (Ogilby, 1839). Wellcome Open Res 9, 368.

Fan, R., Gu, Z., Guang, X., Marin, J.C., Varas, V., Gonzalez, B.A., Wheeler, J.C., Hu, Y., Li, E., Sun, X., Yang, X., Zhang, C., Gao, W., He, J., Munch, K., Corbett-Detig, R., Barbato, M., Pan, S., Zhan, X., Bruford, M.W., Dong, C., 2020. Genomic analysis of the domestication and post-Spanish conquest evolution of the llama and alpaca. Genome Biol 21, 159.

Farre, M., Li, Q., Darolti, I., Zhou, Y., Damas, J., Proskuryakova, A.A., Kulemzina, A.I., Chemnick, L.G., Kim, J., Ryder, O.A., Ma, J., Graphodatsky, A.S., Zhang, G., Larkin, D.M., Lewin, H.A., 2019. An integrated chromosome-scale genome assembly of the Masai giraffe (Giraffa camelopardalis tippelskirchi). Gigascience 8.

Felsenstein, J., 1985. Confidence Limits on Phylogenies: An Approach Using the Bootstrap. Evolution 39, 783–791.

Fitak, R.R., Mohandesan, E., Corander, J., Burger, P.A., 2016. The de novo genome assembly and annotation of a female domestic dromedary of North African origin. Mol Ecol Resour 16, 314–324.

Foote, A., Bunskoek, P., Wellcome Sanger Institute Tree of Life, p., Wellcome Sanger Institute Scientific Operations, D.N.A.P.c., Tree of Life Core Informatics, c., Darwin Tree of Life, C., 2022. The genome sequence of the killer whale, Orcinus orca (Linnaeus, 1758). Wellcome Open Res 7, 250.

Gallus, S., Kumar, V., Bertelsen, M.F., Janke, A., Nilsson, M.A., 2015. A genome survey sequencing of the Java mouse deer (Tragulus javanicus) adds new aspects to the evolution of lineage specific retrotransposons in Ruminantia (Cetartiodactyla). Gene 571, 271–278.

Gao, X., Wang, S., Wang, Y.F., Li, S., Wu, S.X., Yan, R.G., Zhang, Y.W., Wan, R.D., He, Z., Song, R.D., Zhao, X.Q., Wu, D.D., Yang, Q.E., 2022. Long read genome assemblies complemented by single cell RNA-sequencing reveal genetic and cellular mechanisms underlying the adaptive evolution of yak. Nat Commun 13, 4887.

Garcia-Erill, G., Jorgensen, C.H.F., Muwanika, V.B., Wang, X., Rasmussen, M.S., de Jong, Y.A., Gaubert, P., Olayemi, A., Salmona, J., Butynski, T.M., Bertola, L.D., Siegismund, H.R., Albrechtsen, A., Heller, R., 2022. Warthog Genomes Resolve an Evolutionary Conundrum and Reveal Introgression of Disease Resistance Genes. Mol. Biol. Evol. 39.

Groenen, M.A., Archibald, A.L., Uenishi, H., Tuggle, C.K., Takeuchi, Y., Rothschild, M.F., Rogel-Gaillard, C., Park, C., Milan, D., Megens, H.J., Li, S., Larkin, D.M., Kim, H., Frantz, L.A., Caccamo, M., Ahn, H., Aken, B.L., Anselmo, A., Anthon, C., Auvil, L., Badaoui, B., Beattie, C.W., Bendixen, C., Berman, D., Blecha, F., Blomberg, J., Bolund, L., Bosse, M., Botti, S., Bujie, Z., Bystrom, M., Capitanu, B., Carvalho-Silva, D., Chardon, P., Chen, C., Cheng, R., Choi, S.H., Chow, W., Clark, R.C., Clee, C., Crooijmans, R.P., Dawson, H.D., Dehais, P., De Sapio, F., Dibbits, B., Drou, N., Du, Z.Q., Eversole, K., Fadista, J., Fairley, S., Faraut, T., Faulkner, G.J., Fowler, K.E., Fredholm, M., Fritz, E., Gilbert, J.G., Giuffra, E., Gorodkin, J., Griffin, D.K., Harrow, J.L., Hayward, A., Howe, K., Hu, Z.L., Humphray, S.J., Hunt, T., Hornshoj, H., Jeon, J.T., Jern, P., Jones, M., Jurka, J., Kanamori, H., Kapetanovic, R., Kim, J., Kim, J.H., Kim, K.W., Kim, T.H., Larson, G., Lee, K., Lee, K.T., Leggett, R., Lewin, H.A., Li, Y., Liu, W., Loveland, J.E., Lu, Y., Lunney, J.K., Ma, J., Madsen, O., Mann, K., Matthews, L., McLaren, S., Morozumi, T., Murtaugh, M.P., Narayan, J., Nguyen, D.T., Ni, P., Oh, S.J., Onteru, S., Panitz, F., Park, E.W., Park, H.S., Pascal, G., Paudel, Y., Perez-Enciso, M., Ramirez-Gonzalez, R., Reecy, J.M., Rodriguez-Zas, S., Rohrer, G.A., Rund, L., Sang, Y., Schachtschneider, K., Schraiber, J.G., Schwartz, J., Scobie, L., Scott, C., Searle, S., Servin, B., Southey, B.R., Sperber, G., Stadler, P., Sweedler, J.V., Tafer, H., Thomsen, B., Wali, R., Wang, J., Wang, J., White, S., Xu, X., Yerle, M., Zhang, G., Zhang, J., Zhang, J., Zhao, S., Rogers, J., Churcher, C., Schook, L.B., 2012. Analyses of pig genomes provide insight into porcine demography and evolution. Nature 491, 393–398.

Hara-Chikuma, M., Verkman, A.S., 2006. Physiological roles of glycerol-transporting aquaporins: the aquaglyceroporins. Cell. Mol. Life Sci. 63, 1386–1392.

Hidaka, S., Nagashima, A., Kato, A., 2026. Genome-wide identification and solute selectivity of aquaporins in the sharpnose sevengill shark, Heptranchias perlo. Physiol Rep 14, e70895.

Humble, E., Dobrynin, P., Senn, H., Chuven, J., Scott, A.F., Mohr, D.W., Dudchenko, O., Omer, A.D., Colaric, Z., Lieberman Aiden, E., Al Dhaheri, S.S., Wildt, D., Oliaji, S., Tamazian, G., Pukazhenthi, B., Ogden, R., Koepfli, K.P., 2020. Chromosomal-level genome assembly of the scimitar-horned oryx: Insights into diversity and demography of a species extinct in the wild. Mol Ecol Resour 20, 1668–1681.

Imaizumi, G., Ushio, K., Nishihara, H., Braasch, I., Watanabe, E., Kumagai, S., Furuta, T., Matsuzaki, K., Romero, M.F., Kato, A., Nagashima, A., 2024. Functional Divergence in Solute Permeability between Ray-Finned Fish-Specific Paralogs of aqp10. Genome Biol Evol 16.

Ishibashi, K., Imai, M., Sasaki, S., 2000. Cellular localization of aquaporin 7 in the rat kidney. Exp. Nephrol. 8, 252–257.

Ishibashi, K., Morinaga, T., Kuwahara, M., Sasaki, S., Imai, M., 2002. Cloning and identification of a new member of water channel (AQP10) as an aquaglyceroporin. Biochim. Biophys. Acta 1576, 335–340.

Johnson, M., Zaretskaya, I., Raytselis, Y., Merezhuk, Y., McGinnis, S., Madden, T.L., 2008. NCBI BLAST: a better web interface. Nucleic Acids Res 36, W5–9.

Jones, D.T., Taylor, W.R., Thornton, J.M., 1992. The rapid generation of mutation data matrices from protein sequences. Comput. Appl. Biosci. 8, 275–282.

Jones, S.J.M., Taylor, G.A., Chan, S., Warren, R.L., Hammond, S.A., Bilobram, S., Mordecai, G., Suttle, C.A., Miller, K.M., Schulze, A., Chan, A.M., Jones, S.J., Tse, K., Li, I., Cheung, D., Mungall, K.L., Choo, C., Ally, A., Dhalla, N., Tam, A.K.Y., Troussard, A., Kirk, H., Pandoh, P., Paulino, D., Coope, R.J.N., Mungall, A.J., Moore, R., Zhao, Y., Birol, I., Ma, Y., Marra, M., Haulena, M., 2017. The Genome of the Beluga Whale (Delphinapterus leucas). Genes (Basel) 8.

Kato, A., Pipil, S., Ota, C., Kusakabe, M., Watanabe, T., Nagashima, A., Chen, A.P., Islam, Z., Hayashi, N., Wong, M.K., Komada, M., Romero, M.F., Takei, Y., 2024. Convergent gene losses and pseudogenizations in multiple lineages of stomachless fishes. Commun Biol 7, 408.

Koyama, Y., Yamamoto, T., Tani, T., Nihei, K., Kondo, D., Funaki, H., Yaoita, E., Kawasaki, K., Sato, N., Hatakeyama, K., Kihara, I., 1999. Expression and localization of aquaporins in rat gastrointestinal tract. Am. J. Physiol. 276, C621–627.

Kumar, S., Stecher, G., Suleski, M., Sanderford, M., Sharma, S., Tamura, K., 2024. MEGA12: Molecular Evolutionary Genetic Analysis Version 12 for Adaptive and Green Computing. Mol. Biol. Evol. 41.

Kumar, S., Suleski, M., Craig, J.M., Kasprowicz, A.E., Sanderford, M., Li, M., Stecher, G., Hedges, S.B., 2022. TimeTree 5: An Expanded Resource for Species Divergence Times. Mol. Biol. Evol. 39.

Laforenza, U., 2012. Water channel proteins in the gastrointestinal tract. Mol. Aspects Med. 33, 642–650.

Lamb, S., Taylor, A.M., Hughes, T.A., McMillan, B.R., Larsen, R.T., Khan, R., Weisz, D., Dudchenko, O., Aiden, E.L., Edelman, N.B., Frandsen, P.B., 2021. De novo chromosome-length assembly of the mule deer (Odocoileus hemionus) genome. GigaByte 2021, gigabyte34.

Lee, K., Kim, H.K., Park, S.K., Sohn, H., Cho, Y., Choi, Y.M., Jeong, D.G., Kim, J.H., 2018. First report of the occurrence and whole-genome characterization of Edwardsiella tarda in the false killer whale (Pseudorca crassidens). J. Vet. Med. Sci. 80, 1041–1046.

Li, K., Cappelletti, E., Dessaix, C., Ciosek, J., Robyn, E., Johnson, L., AbouEl Ela, N.H., Arias, X., Adelson, D.L., Raudsepp, T., Piras, F.M., Smith, M.L., Hudson, E., Pickett, B.D., Koren, S., Walenz, B.P., Brooks, S.Y., Sison, C., Crawford, J., Bouffard, G., Phillippy, A.M., Miller, D., Antczak, D.F., Cullen, J., Stroupe, S., Davis, B.W., McCue, M., Durward-Akhurst, S.A., Petersen, J.L., Giulotto, E., Kalbfleisch, T., 2026. Phased T2T horse and donkey assemblies from a mule reveal peculiar equid centromere evolution. Cell Genom 6, 101307.

Liu, C., Gao, J., Cui, X., Li, Z., Chen, L., Yuan, Y., Zhang, Y., Mei, L., Zhao, L., Cai, D., Hu, M., Zhou, B., Li, Z., Qin, T., Si, H., Li, G., Lin, Z., Xu, Y., Zhu, C., Yin, Y., Zhang, C., Xu, W., Li, Q., Wang, K., Gilbert, M.T.P., Heller, R., Wang, W., Huang, J., Qiu, Q., 2021. A towering genome: Experimentally validated adaptations to high blood pressure and extreme stature in the giraffe. Sci Adv 7.

Login, F.H., Nejsum, L.N., 2023. Aquaporin water channels: roles beyond renal water handling. Nat Rev Nephrol 19, 604–618.

London, E.W., Roca, A.L., Novakofski, J.E., Mateus-Pinilla, N.E., 2022. A De Novo Chromosome-Level Genome Assembly of the White-Tailed Deer, Odocoileus Virginianus. J. Hered. 113, 479–489.

Madeira, F., Madhusoodanan, N., Lee, J., Eusebi, A., Niewielska, A., Tivey, A.R.N., Lopez, R., Butcher, S., 2024. The EMBL-EBI Job Dispatcher sequence analysis tools framework in 2024. Nucleic Acids Res 52, W521–W525.

Martin, F.J., Amode, M.R., Aneja, A., Austine-Orimoloye, O., Azov, A.G., Barnes, I., Becker, A., Bennett, R., Berry, A., Bhai, J., Bhurji, S.K., Bignell, A., Boddu, S., Branco Lins, P.R., Brooks, L., Ramaraju, S.B., Charkhchi, M., Cockburn, A., Da Rin Fiorretto, L., Davidson, C., Dodiya, K., Donaldson, S., El Houdaigui, B., El Naboulsi, T., Fatima, R., Giron, C.G., Genez, T., Ghattaoraya, G.S., Martinez, J.G., Guijarro, C., Hardy, M., Hollis, Z., Hourlier, T., Hunt, T., Kay, M., Kaykala, V., Le, T., Lemos, D., Marques-Coelho, D., Marugan, J.C., Merino, G.A., Mirabueno, L.P., Mushtaq, A., Hossain, S.N., Ogeh, D.N., Sakthivel, M.P., Parker, A., Perry, M., Pilizota, I., Prosovetskaia, I., Perez-Silva, J.G., Salam, A.I.A., Saraiva-Agostinho, N., Schuilenburg, H., Sheppard, D., Sinha, S., Sipos, B., Stark, W., Steed, E., Sukumaran, R., Sumathipala, D., Suner, M.M., Surapaneni, L., Sutinen, K., Szpak, M., Tricomi, F.F., Urbina-Gomez, D., Veidenberg, A., Walsh, T.A., Walts, B., Wass, E., Willhoft, N., Allen, J., Alvarez-Jarreta, J., Chakiachvili, M., Flint, B., Giorgetti, S., Haggerty, L., Ilsley, G.R., Loveland, J.E., Moore, B., Mudge, J.M., Tate, J., Thybert, D., Trevanion, S.J., Winterbottom, A., Frankish, A., Hunt, S.E., Ruffier, M., Cunningham, F., Dyer, S., Finn, R.D., Howe, K.L., Harrison, P.W., Yates, A.D., Flicek, P., 2023. Ensembl 2023. Nucleic Acids Res 51, D933–D941.

Martinez-Viaud, K.A., Lawley, C.T., Vergara, M.M., Ben-Zvi, G., Biniashvili, T., Baruch, K., St Leger, J., Le, J., Natarajan, A., Rivera, M., Guillergan, M., Jaeger, E., Steffy, B., Zimin, A., 2019. New de novo assembly of the Atlantic bottlenose dolphin (Tursiops truncatus) improves genome completeness and provides haplotype phasing. Gigascience 8.

Masonbrink, R.E., Alt, D., Bayles, D.O., Boggiatto, P., Edwards, W., Tatum, F., Williams, J., Wilson-Welder, J., Zimin, A., Severin, A., Olsen, S., 2021. A pseudomolecule assembly of the Rocky Mountain elk genome. PLoS One 16, e0249899.

Miller, J.M., Moore, S.S., Stothard, P., Liao, X., Coltman, D.W., 2015. Harnessing cross-species alignment to discover SNPs and generate a draft genome sequence of a bighorn sheep (Ovis canadensis). BMC Genomics 16, 397.

Ming, L., Wang, Z., Yi, L., Batmunkh, M., Liu, T., Siren, D., He, J., Juramt, N., Jambl, T., Li, Y. Jirimutu, 2020. Chromosome-level assembly of wild Bactrian camel genome reveals organization of immune gene loci. Mol Ecol Resour 20.

Mobasheri, A., Shakibaei, M., Marples, D., 2004. Immunohistochemical localization of aquaporin 10 in the apical membranes of the human ileum: a potential pathway for luminal water and small solute absorption. Histochem. Cell Biol. 121, 463–471.

Morinaga, T., Nakakoshi, M., Hirao, A., Imai, M., Ishibashi, K., 2002. Mouse aquaporin 10 gene (AQP10) is a pseudogene. Biochem. Biophys. Res. Commun. 294, 630–634.

Moskalev Acapital A, c., Kudryavtseva, A.V., Graphodatsky, A.S., Beklemisheva, V.R., Serdyukova, N.A., Krutovsky, K.V., Sharov, V.V., Kulakovskiy, I.V., Lando, A.S., Kasianov, A.S., Kuzmin, D.A., Putintseva, Y.A., Feranchuk, S.I., Shaposhnikov, M.V., Fraifeld, V.E., Toren, D., Snezhkina, A.V., Sitnik, V.V., 2017. De novo assembling and primary analysis of genome and transcriptome of gray whale Eschrichtius robustus. BMC Evol Biol 17, 258.

Motoshima, T., Nagashima, A., Ota, C., Oka, H., Hosono, K., Braasch, I., Nishihara, H., Kato, A., 2023. Na+-Cl- cotransporter 2 is not fish-specific and is widely found in amphibians, non-avian reptiles, and select mammals. Physiol Genomics.

Nagashima, A., Hidaka, S., Ota, C., Yamanaka, D., Ito, K., Nakada, T., Furuta, T., Kato, A., 2025a. Loss and Gain of Aqp10 Paralogs With Broad Solute Selectivity in Anguillid Eels. Genome Biol Evol 17.

Nagashima, A., Kato, A., 2026. Elasmobranch Aqp10 paralogs differ in glycerol permeability. Comp. Biochem. Physiol. B. Biochem. Mol. Biol. 282, 111191.

Nagashima, A., Nagai, N., Ota, C., Ushio, K., Kato, A., 2025b. Retention and pseudogenization of aquaporin-10 in Rodentia. Biochem. Biophys. Res. Commun. 756, 151608.

Nagashima, A., Ushio, K., Nishihara, H., Akimoto, J., Kato, A., Furuta, T., 2025c. Aquaporin 10 paralogs exhibit evolutionarily altered urea and boric acid permeabilities based on the amino acid residues at positions 1 and 3 in the ar/R region. Am. J. Physiol. Regul. Integr. Comp. Physiol. 329, R423–R436.

Nei, M., Gojobori, T., 1986. Simple methods for estimating the numbers of synonymous and nonsynonymous nucleotide substitutions. Mol. Biol. Evol. 3, 418–426.

Oppenheimer, J., Rosen, B.D., Heaton, M.P., Vander Ley, B.L., Shafer, W.R., Schuetze, F.T., Stroud, B., Kuehn, L.A., McClure, J.C., Barfield, J.P., Blackburn, H.D., Kalbfleisch, T.S., Bickhart, D.M., Davenport, K.M., Kuhn, K.L., Green, R.E., Shapiro, B., Smith, T.P.L., 2021. A Reference Genome Assembly of American Bison, Bison bison bison. J. Hered. 112, 174–183.

Pink, R.C., Wicks, K., Caley, D.P., Punch, E.K., Jacobs, L., Carter, D.R., 2011. Pseudogenes: pseudo-functional or key regulators in health and disease? RNA 17, 792–798.

Prewer, E., Kutz, S., Leclerc, L.M., Kyle, C.J., 2022. Draft Genome Assembly of an Iconic Arctic Species: Muskox (Ovibos moschatus). Genes (Basel) 13.

Qiao, G., Xu, P., Guo, T., Wu, Y., Lu, X., Zhang, Q., He, X., Zhu, S., Zhao, H., Lei, Z., Sun, W., Yang, B., Yue, Y., 2022. Genetic Basis of Dorper Sheep (Ovis aries) Revealed by Long-Read De Novo Genome Assembly. Front Genet 13, 846449.

Rajput, S., Gautam, D., Vats, A., Roshan, M., Goyal, P., Rana, C., S, M.p., Ludri, A., De, S., 2024. Aquaporin (AQP) gene family in Buffalo and Goat: Molecular characterization and their expression analysis. Int. J.Biol. Macromol. 280, 136145.

Rangwala, S.H., Kuznetsov, A., Ananiev, V., Asztalos, A., Borodin, E., Evgeniev, V., Joukov, V., Lotov, V., Pannu, R., Rudnev, D., Shkeda, A., Weitz, E.M., Schneider, V.A., 2021. Accessing NCBI data using the NCBI Sequence Viewer and Genome Data Viewer (GDV). Genome Res. 31, 159–169.

Rhie, A., McCarthy, S.A., Fedrigo, O., Damas, J., Formenti, G., Koren, S., Uliano-Silva, M., Chow, W., Fungtammasan, A., Kim, J., Lee, C., Ko, B.J., Chaisson, M., Gedman, G.L., Cantin, L.J., Thibaud-Nissen, F., Haggerty, L., Bista, I., Smith, M., Haase, B., Mountcastle, J., Winkler, S., Paez, S., Howard, J., Vernes, S.C., Lama, T.M., Grutzner, F., Warren, W.C., Balakrishnan, C.N., Burt, D., George, J.M., Biegler, M.T., Iorns, D., Digby, A., Eason, D., Robertson, B., Edwards, T., Wilkinson, M., Turner, G., Meyer, A., Kautt, A.F., Franchini, P., Detrich, H.W., 3rd, Svardal, H., Wagner, M., Naylor, G.J.P., Pippel, M., Malinsky, M., Mooney, M., Simbirsky, M., Hannigan, B.T., Pesout, T., Houck, M., Misuraca, A., Kingan, S.B., Hall, R., Kronenberg, Z., Sovic, I., Dunn, C., Ning, Z., Hastie, A., Lee, J., Selvaraj, S., Green, R.E., Putnam, N.H., Gut, I., Ghurye, J., Garrison, E., Sims, Y., Collins, J., Pelan, S., Torrance, J., Tracey, A., Wood, J., Dagnew, R.E., Guan, D., London, S.E., Clayton, D.F., Mello, C.V., Friedrich, S.R., Lovell, P.V., Osipova, E., Al-Ajli, F.O., Secomandi, S., Kim, H., Theofanopoulou, C., Hiller, M., Zhou, Y., Harris, R.S., Makova, K.D., Medvedev, P., Hoffman, J., Masterson, P., Clark, K., Martin, F., Howe, K., Flicek, P., Walenz, B.P., Kwak, W., Clawson, H., Diekhans, M., Nassar, L., Paten, B., Kraus, R.H.S., Crawford, A.J., Gilbert, M.T.P., Zhang, G., Venkatesh, B., Murphy, R.W., Koepfli, K.P., Shapiro, B., Johnson, W.E., Di Palma, F., Marques-Bonet, T., Teeling, E.C., Warnow, T., Graves, J.M., Ryder, O.A., Haussler, D., O’Brien, S.J., Korlach, J., Lewin, H.A., Howe, K., Myers, E.W., Durbin, R., Phillippy, A.M., Jarvis, E.D., 2021. Towards complete and error-free genome assemblies of all vertebrate species. Nature 592, 737–746.

Richardson, M.F., Munyard, K., Croft, L.J., Allnutt, T.R., Jackling, F., Alshanbari, F., Jevit, M., Wright, G.A., Cransberg, R., Tibary, A., Perelman, P., Appleton, B., Raudsepp, T., 2019. Chromosome-Level Alpaca Reference Genome VicPac3.1 Improves Genomic Insight Into the Biology of New World Camelids. Front Genet 10, 586.

Rojek, A., Praetorius, J., Frokiaer, J., Nielsen, S., Fenton, R.A., 2008. A current view of the mammalian aquaglyceroporins. Annu. Rev. Physiol. 70, 301–327.

Russell, J.B., Muck, R.E., Weimer, P.J., 2009. Quantitative analysis of cellulose degradation and growth of cellulolytic bacteria in the rumen. FEMS Microbiol Ecol 67, 183–197.

Saitou, N., Nei, M., 1987. The neighbor-joining method: a new method for reconstructing phylogenetic trees. Mol. Biol. Evol. 4, 406–425.

Si, J., Dai, D., Gorkhali, N.A., Wang, M., Wang, S., Sapkota, S., Kadel, R.C., Sadaula, A., Dhakal, A., Faruque, M.O., Omar, A.I., Sari, E.M., Ashari, H., Dagong, M.I.A., Yindee, M., Rushdi, H.E., Elregalaty, H., Amin, A., Radwan, M.A., Pham, L.D., Hulugalla, W., Silva, G., Zheng, W., Mansoor, S., Ali, M.B., Vahidi, F., Al-Bayatti, S.A., Pauciullo, A., Lenstra, J.A., Barker, J.S.F., Fang, L., Wu, D.D., Han, J., Zhang, Y., 2024. Complete Genomic Landscape Reveals Hidden Evolutionary History and Selection Signature in Asian Water Buffaloes (Bubalus bubalis). Adv Sci (Weinh), e2407615.

Sohara, E., Rai, T., Sasaki, S., Uchida, S., 2006. Physiological roles of AQP7 in the kidney: Lessons from AQP7 knockout mice. Biochim. Biophys. Acta 1758, 1106–1110.

Springer, M.S., Guerrero-Juarez, C.F., Huelsmann, M., Collin, M.A., Danil, K., McGowen, M.R., Oh, J.W., Ramos, R., Hiller, M., Plikus, M.V., Gatesy, J., 2021. Genomic and anatomical comparisons of skin support independent adaptation to life in water by cetaceans and hippos. Curr. Biol. 31, 2124–2139 e2123.

Stewart, G.S., Smith, C.P., 2005. Urea nitrogen salvage mechanisms and their relevance to ruminants, non-ruminants and man. Nutr Res Rev 18, 49–62.

Tanaka, Y., Morishita, Y., Ishibashi, K., 2015. Aquaporin10 is a pseudogene in cattle and their relatives. Biochem Biophys Rep 1, 16–21.

Tran, Y.H., Xu, Z., Kato, A., Mistry, A.C., Goya, Y., Taira, M., Brandt, S.J., Hirose, S., 2006. Spliced isoforms of LIM-domain-binding protein (CLIM/NLI/Ldb) lacking the LIM-interaction domain. J Biochem 140, 105–119.

Tutar, Y., 2012. Pseudogenes. Comp Funct Genomics 2012, 424526.

Weldenegodguad, M., Pokharel, K., Ming, Y., Honkatukia, M., Peippo, J., Reilas, T., Roed, K.H., Kantanen, J., 2020. Genome sequence and comparative analysis of reindeer (Rangifer tarandus) in northern Eurasia. Sci Rep 10, 8980.

Winter, S., Coimbra, R.T.F., Helsen, P., Janke, A., 2022. A Chromosome-Scale Genome Assembly of the Okapi (Okapia johnstoni). J. Hered. 113, 568–576.

Xu, B., Chen, J., Song, P., Gu, H., Jiang, F., Li, B., Wei, Q., Zhang, T., 2024. A high-quality chromosome-level reference genome assembly of Tibetan antelope (Pantholops hodgsonii). Sci Data 11, 1215.

Yi, L., Dalai, M., Su, R., Lin, W., Erdenedalai, M., Luvsantseren, B., Chimedtseren, C., Wang, Z., Hasi, S., 2020. Whole-genome sequencing of wild Siberian musk deer (Moschus moschiferus) provides insights into its genetic features. BMC Genomics 21, 108.

Yilmaz, O., Chauvigne, F., Ferre, A., Nilsen, F., Fjelldal, P.G., Cerda, J., Finn, R.N., 2020. Unravelling the Complex Duplication History of Deuterostome Glycerol Transporters. Cells 9.

Yim, H.S., Cho, Y.S., Guang, X., Kang, S.G., Jeong, J.Y., Cha, S.S., Oh, H.M., Lee, J.H., Yang, E.C., Kwon, K.K., Kim, Y.J., Kim, T.W., Kim, W., Jeon, J.H., Kim, S.J., Choi, D.H., Jho, S., Kim, H.M., Ko, J., Kim, H., Shin, Y.A., Jung, H.J., Zheng, Y., Wang, Z., Chen, Y., Chen, M., Jiang, A., Li, E., Zhang, S., Hou, H., Kim, T.H., Yu, L., Liu, S., Ahn, K., Cooper, J., Park, S.G., Hong, C.P., Jin, W., Kim, H.S., Park, C., Lee, K., Chun, S., Morin, P.A., O’Brien, S.J., Lee, H., Kimura, J., Moon, D.Y., Manica, A., Edwards, J., Kim, B.C., Kim, S., Wang, J., Bhak, J., Lee, H.S., Lee, J.H., 2014. Minke whale genome and aquatic adaptation in cetaceans. Nat. Genet. 46, 88–92.

Zhang, Z.D., Frankish, A., Hunt, T., Harrow, J., Gerstein, M., 2010. Identification and analysis of unitary pseudogenes: historic and contemporary gene losses in humans and other primates. Genome Biol 11, R26.

Zhou, C., Peng, K., Liu, Y., Zhang, R., Zheng, X., Yue, B., Du, C., Wu, Y., 2023. Comparative Analyses Reveal the Genetic Mechanism of Ambergris Production in the Sperm Whale Based on the Chromosome-Level Genome. Animals (Basel) 13.

Zhou, X., Sun, F., Xu, S., Fan, G., Zhu, K., Liu, X., Chen, Y., Shi, C., Yang, Y., Huang, Z., Chen, J., Hou, H., Guo, X., Chen, W., Chen, Y., Wang, X., Lv, T., Yang, D., Zhou, J., Huang, B., Wang, Z., Zhao, W., Tian, R., Xiong, Z., Xu, J., Liang, X., Chen, B., Liu, W., Wang, J., Pan, S., Fang, X., Li, M., Wei, F., Xu, X., Zhou, K., Wang, J., Yang, G., 2013. Baiji genomes reveal low genetic variability and new insights into secondary aquatic adaptations. Nat Commun 4, 2708.

Zimin, A.V., Delcher, A.L., Florea, L., Kelley, D.R., Schatz, M.C., Puiu, D., Hanrahan, F., Pertea, G., Van Tassell, C.P., Sonstegard, T.S., Marcais, G., Roberts, M., Subramanian, P., Yorke, J.A., Salzberg, S.L., 2009. A whole-genome assembly of the domestic cow, Bos taurus. Genome Biol 10, R42.

Zoonomia, C., 2020. A comparative genomics multitool for scientific discovery and conservation. Nature 587, 240–245.

Zurano, J.P., Magalhaes, F.M., Asato, A.E., Silva, G., Bidau, C.J., Mesquita, D.O., Costa, G.C., 2019. Cetartiodactyla: Updating a time-calibrated molecular phylogeny. Mol. Phylogenet. Evol. 133, 256–262.

