## Supplementary material for "Evolutionary history of aquaporin-10 pseudogenization in Cetartiodactyla": Fig. S1

**A** **exon 3-6**

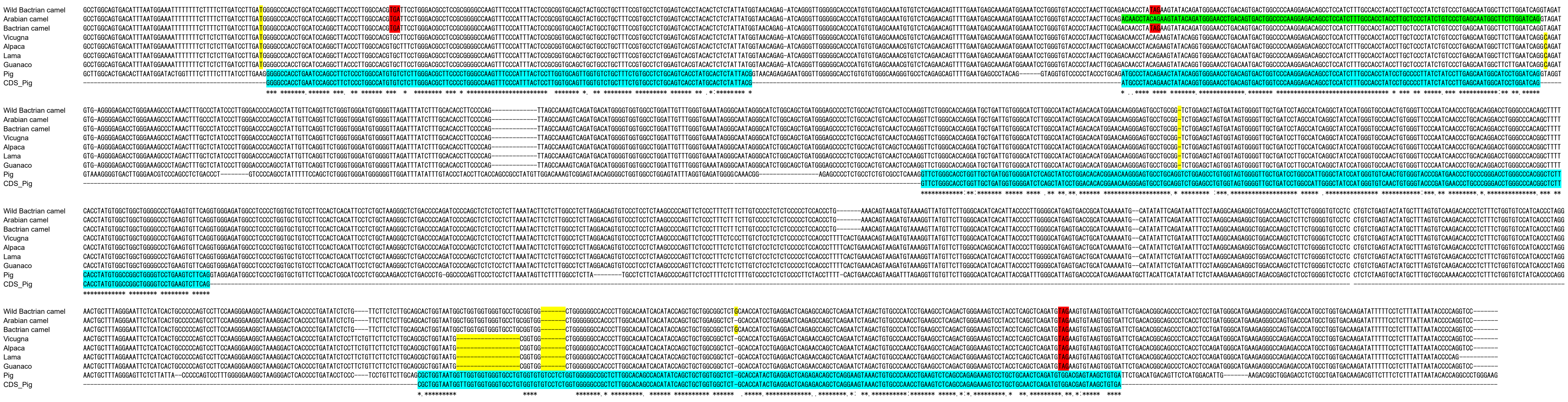


**B**

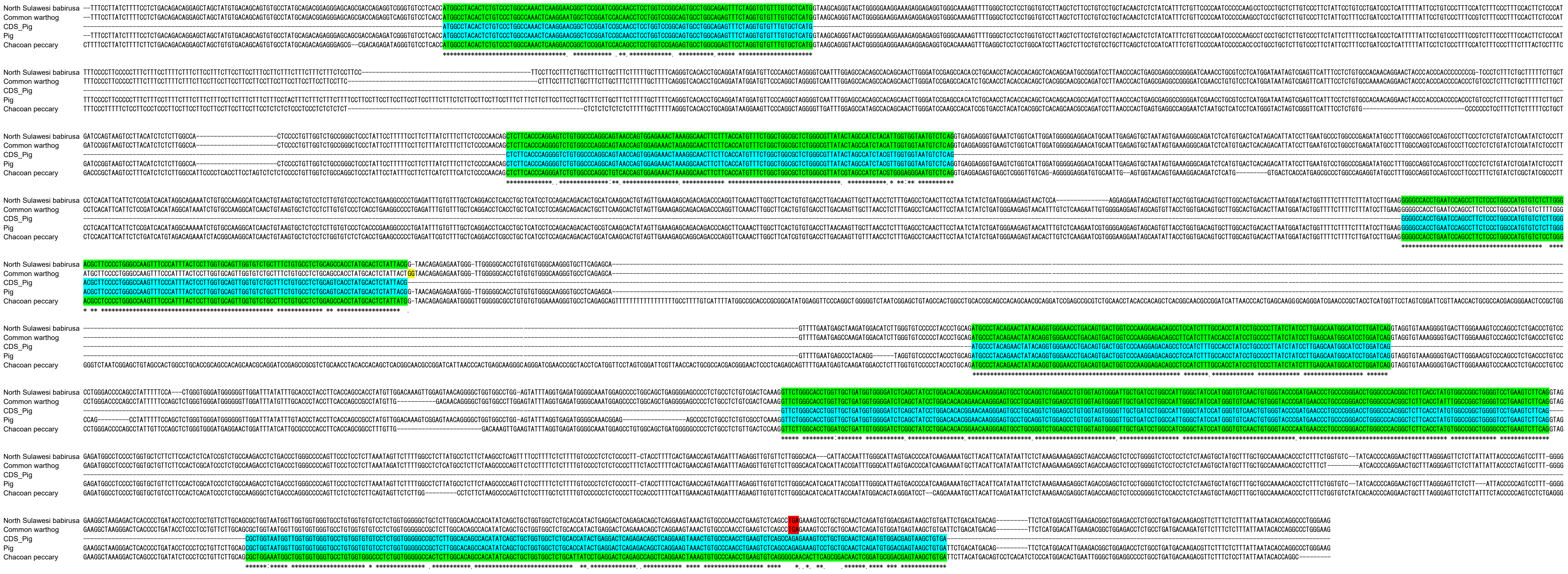


以下 pronghorn 以外

**C** exon 1

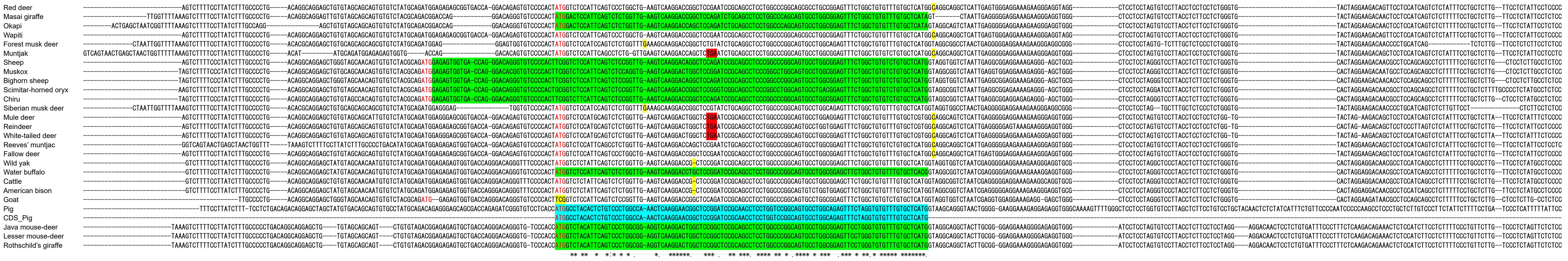


**D** exon 2-6

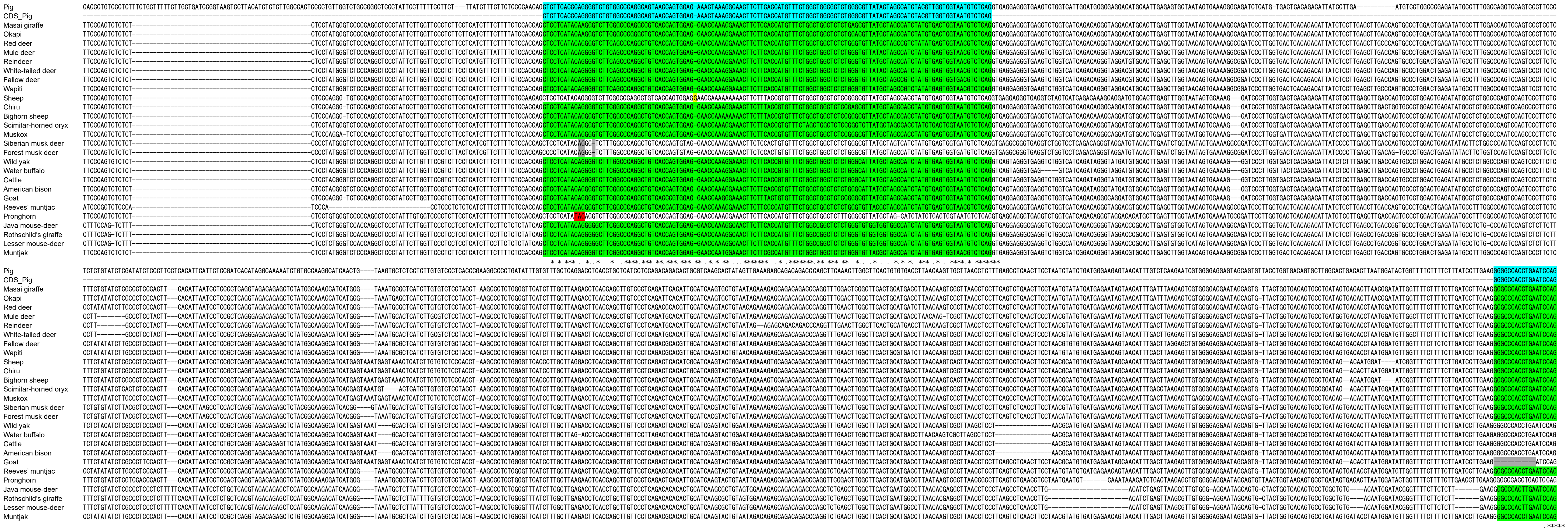

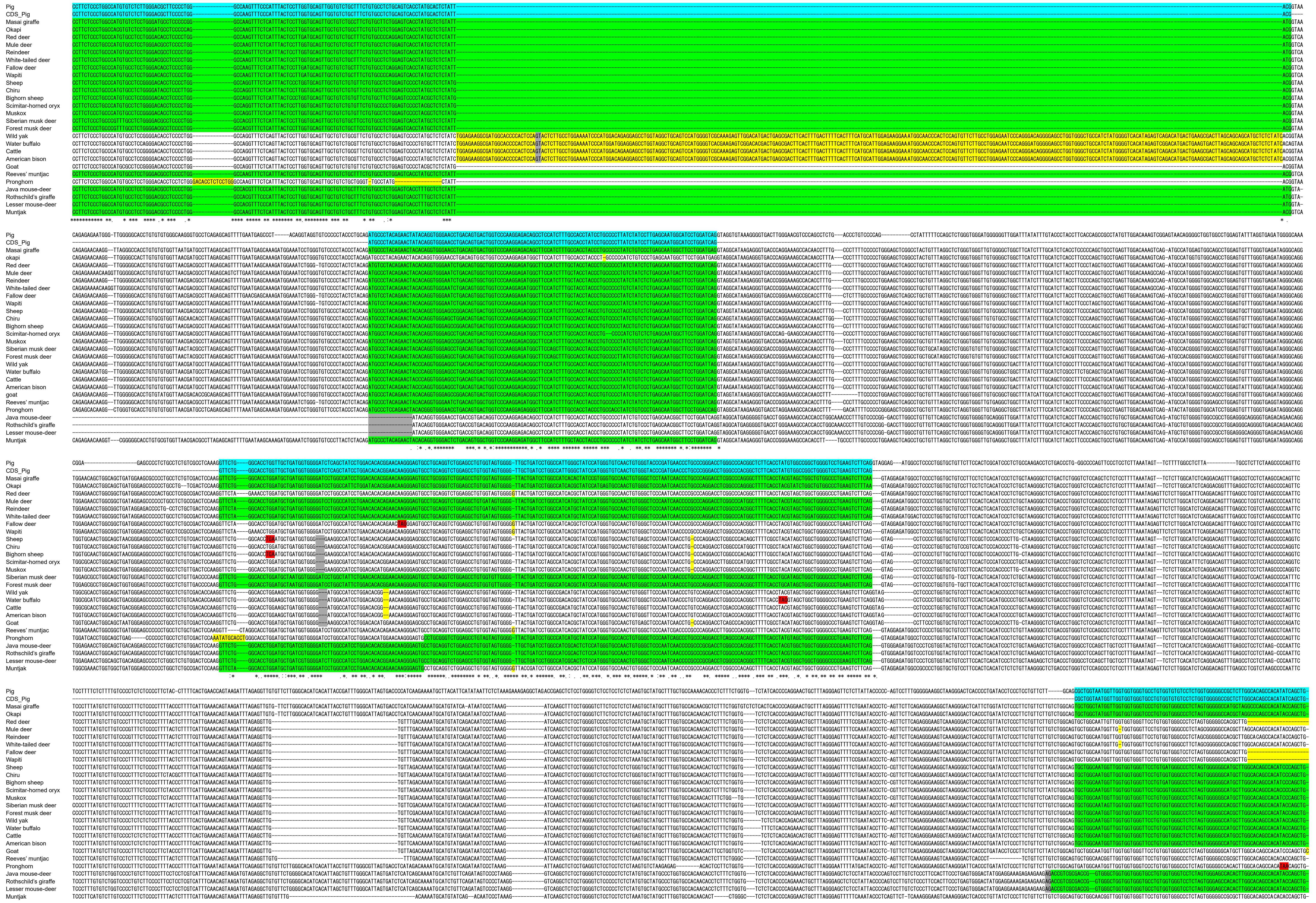
Pig
CDS_Pig
Masai giraffe
Okapi
Red deer
Mule deer
Reindeer
White-tailed deer
Fallow deer
Wapiti
Sheep
Chiru
Bighorn sheep
Scimitar-horned oryx
Muskox
Siberian musk deer
Forest musk deer
Wild yak
Water buffalo
Cattle
American bison
Goat
Reeves' muntjac
Pronghorn
Java mouse-deer
Rothschild's giraffe
Lesser mouse-deer
Muntjak

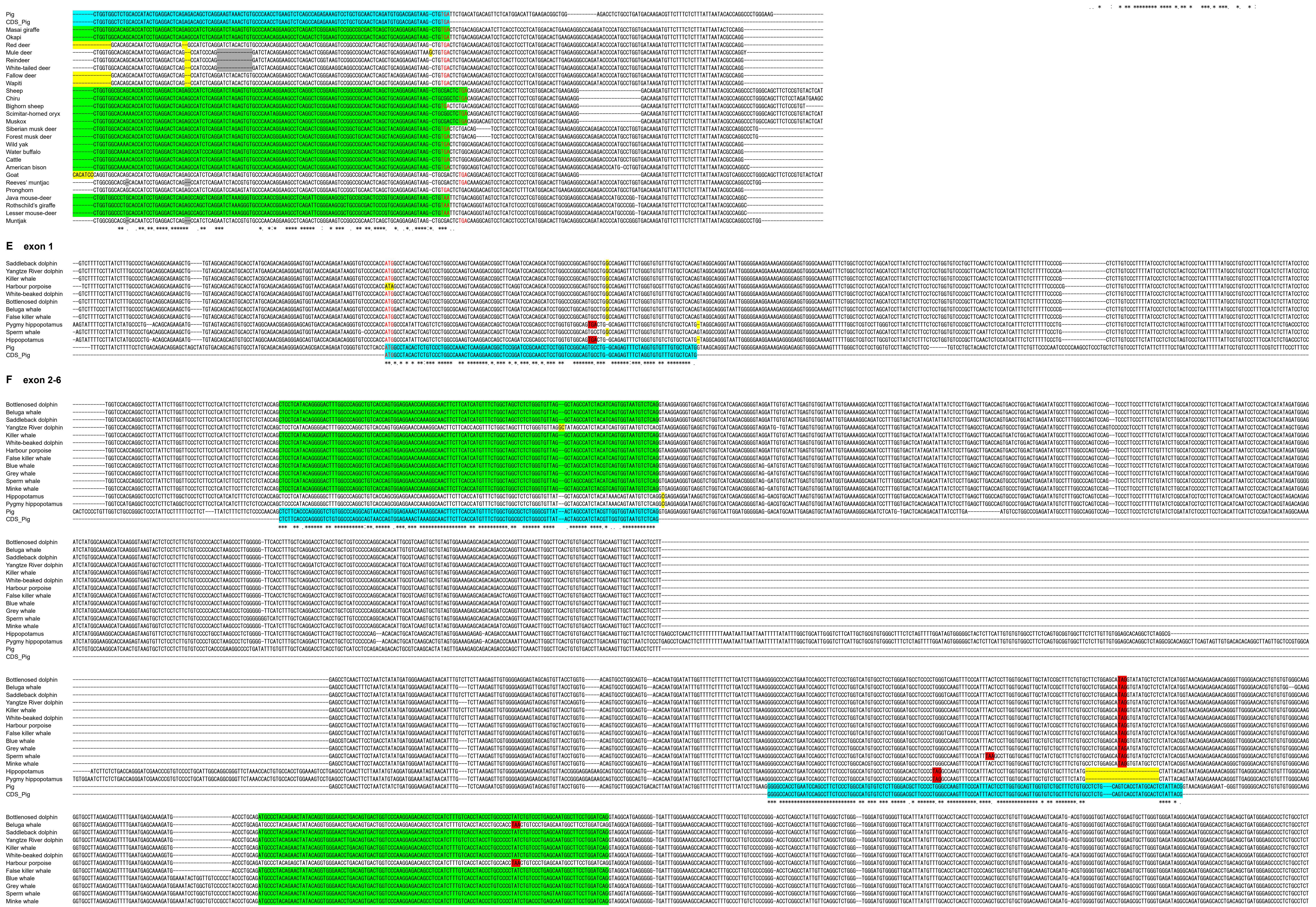
Pig
CDS_Pig
Masai giraffe
Okapi
Red deer
Mule deer
Reindeer
White-tailed deer
Fallow deer
Wapiti
Sheep
Chiru
Bighorn sheep
Scimitar-horned oryx
Muskox
Siberian musk deer
Forest musk deer
Wild yak
Water buffalo
Cattle
American bison
Goat
Reeves' muntjac
Pronghorn
Java mouse-deer
Rothschild's giraffe
Lesser mouse-deer
Muntjak
E exon 1
Saddleback dolphin
Yangtze River dolphin
Killer whale
Harbour porpoise
White-beaked dolphin
Bottlenosed dolphin
Beluga whale
False killer whale
Pygmy hippopotamus
Sperm whale
Hippopotamus
Pig
CDS_Pig
F exon 2-6
Bottlenose dolphin
Beluga whale
Saddleback dolphin
Yangtze River dolphin
Killer whale
White-beaked dolphin
Harbour porpoise
False killer whale
Blue whale
Grey whale
Sperm whale
Minke whale
Hippopotamus
Pygmy hippopotamus
Pig
CDS_Pig

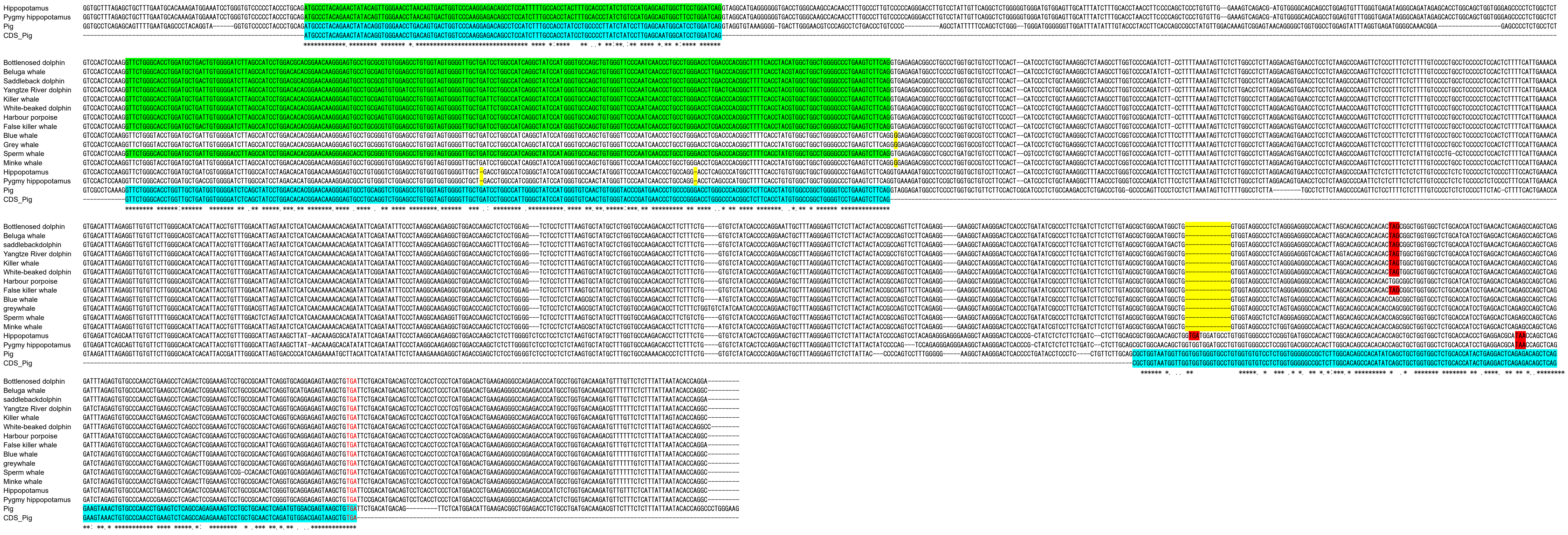

**Fig. S1.** Multiple alignment of *Aqp10* pseudogenes (*Aqp10p*).
The nucleotide sequences of six exons for *Aqp10p* were aligned with Pig and/or *Aqp10*. Intact exons and CDSs of Pig *Aqp10* are highlighted in light blue. Intact exons of *Aqp10p* are highlighted in green, and stop codons are highlighted in red. Frameshifts, non-canonical splice sites, and mutations in the start codon are highlighted in yellow. Poly(N) region between exon 3 and exon 4 of Chacoan peccary were deleted.
