## Supplementary material for "Evolutionary history of aquaporin-10 pseudogenization in Cetartiodactyla": Table S3

**Table S3.** Analysis of the synteny of *Aqp10* in the Cetartiodactyla and horse genome databases

| No. | Common name | Species | Genome Database | synteny | synteny | synteny | synteny | synteny | synteny |
|---|---|---|---|---|---|---|---|---|---|
| 1 | Pig | Sus scrofa | GCA_054392235.2 | Chr. 4 (minus) | >ubap2l XM_013997060.2 | >hax1 XM_021089792.1 | >aqp10 NM_001128454.1 | >atp8b2 XM_005663380.3 | >il6r NM_214403.1 |
| 2 | Common warthog | Phacochoerus africanus | GCA_016906955.1 | Chr. 6 (minus) | >ubap2l XM_047784469.1 | >hax1 XM_047784507.1 | >aqp10 XM_047782895.1 | >atp8b2 XM_047784467.1 | >il6r XM_047784465.1 |
| 3 | Babyrousa | Babyrousa celebensis | GCA_057927545.1 | Chr. 8 (minus)JAPYYC010000008.1 | >ubap2l region: 82514532-82473963 | >hax1 region:82471504-82469695 | >aqp10 region:82457166-82454240 | >atp8b2 region:82449700-82432831 | >il6r region:82373280-82348397 |
| 4 | Chacoan peccary | Catagonus wagneri | GCA_004024745.2 | BS18 Sc2sB16_25;HRSCAF=9013_0 (minus)PVHT020000034.1 | >ubap2l region: 688226-648556 | >hax1 region:646129-644264 | >aqp10 region:635606-631251 | >atp8b2 region:626667-607500 | >il6r region:545216-513765 |
| 5 | Arabian camel | Camelus dromedarius | GCA_036321535.1 | NW_011513412.1 (minus) | >ubap2l XM_045515979.1 | >hax1 XM_045516002.1 | >aqp10 XM_010954085.1 | >atp8b2 XM_045515977.1 | >il6r XM_045515976.1 |
| 6 | Wild bactrian camel | Camelus ferus | GCA_009834535.1 | Chr. 21 (minus) | >ubap2l M_032464163.1 | >hax1 XM_006177220.3 | >aqp10 XM_032464768.1 | >atp8b2 XM_006177226.3 | >il6r XM_032464159.1 |
| 7 | Bactrian camel | Camelus bactrianus | GCA_048773025.1 | NW_011513412.1 (minus) | >ubap2l XM_045515979.1 | >hax1 XM_045516002.1 | >aqp10 XM_010954085.1 | >atp8b2 XM_045515977.1 | >il6r XM_045515976.1 |
| 8 | Vicugna | Vicugna vicugna | GCA_013265495.1 | contig26126 (minus) PNXW01026126.1 | >ubap2l region: 133466-101731 | >hax1 region: 99266-97469 | >aqp10 region: 91504-90057 | >atp8b2 region: 85596-69438 | >il6r region: 23631-18587 |
| 9 | Guanaco | Lama guanicoe cacsilensis | GCA_013239625.1 | contig13028 (minus) PNXV01013028.1 | >ubap2l region: 320317-288573 | >hax1 region: 286117-284319 | >aqp10 region: 278294-276847 | >atp8b2 region: 272295-256031 | >il6r region: 210007-178478 |
| 10 | Lama | Lama glama chaku | GCA_013239585.1 | contig39556 (plus) PNXU01039556.1 | | >hax1 region: 1384-3182 | >aqp10 region: 9439-10886 | | |
| 11 | Alpaca | Vicugna pacos | GCA_048564905.1 | NW_021964201.1 (plus) | >ubap2l XM_031689028.1 | >hax1 region: 2505612-2507397 | >aqp10 XM_006220151.3 | >atp8b2 XM_015248276.2 | >il6r XM_031689061.1 |
| 12 | Java mouse-deer | Tragulus javanicus | GCA_004024965.2 | US108 ScTR3Pl_88;HRSCAF=25073 (plus) PVHZ020000093.1 | >ubap2l region: 2467451-2503099 | >hax1 region: 21661-26455 | >aqp10 region: 2525217-2527843 | >atp8b2 region: 2532236-2550635 | >il6r region: 2618050-2649912 |
| 13 | lesser mouse-deer | Tragulus kanchil | GCA_022376925.1 | LMD-DV tarseq_1137 (plus) SJXW01010930.1 | | | >aqp10 region:45890-48404 | >atp8b2 region:52806-71673 | |
| 14 | Pronghorn | Antilocapra americana | GCA_051176615.2 | contig139147_2 (minus) SRVM01010816.1 | >ubap2l region: 304230-268548 | | >aqp10 region: 250339-248182 | >atp8b2 region: 243654-227365 | >il6r region: 171821-142834 |
| 15 | Masai giraffe | Giraffa tippelskirchi | GCA_054371585.1 | Chr. 5 (minus) RAWU01000023.1 | >ubap2l region: 171387779-171351696 | >hax1 region: 38938-40695 | >aqp10 region: 171333552-171330930 | >atp8b2 region: 171326631-171310310 | >il6r region: 171253796-171221851 |
| 16 | Rothschild's giraffe | Giraffa camelopardalis rothschildi | GCA_017591445.1 | scaffold4609b103139e642173_34 (plus) LVKQ01130786.1 | >ubap2l region: 432-36488 | >hax1 region: - | >aqp10 region: 55136-57758 | >atp8b2 region:62321-78642 | |
| 17 | Okapi | Okapia johnstoni | GCA_024291935.2 | scaffold13182b47170e177359_8 (plus) LVCL010199069.1 | | | >aqp10 region: 6722-8895 | >atp8b2 region: 13454-29780 | |
| 18 | Wapiti | Cervus canadensis | GCA_019320065.1 | Chr. 2 (minus) | >ubap2l XM_043455468.1 | | >aqp10 XM_043455414.1 | >atp8b2 XM_043455375.1 | >il6r XM_043455362.1 |
| 19 | Red deer | Cervus elaphus | GCA_910594005.1 | Chr. 20 (minus) | >ubap2l XM_043875642.1 | >hax1 XM_061120708.1 | >aqp10 XM_043876762.1 | >atp8b2 XM_043876758.1 | >il6r XM_043877211.1 |
| 20 | Fallow deer | Dama dama | GCA_033118175.2 | Chr. 20 (minus) | >ubap2l XM_061120711.1 | >hax1 region: 566447-564679 | >aqp10 XM_061119869.1 | >atp8b2 XM_061120706.1 | >il6r XM_061120705.1 |

| | | | | | | | | | |
|---|---|---|---|---|---|---|---|---|---|
| 21 | Muntjak | Muntiacus muntjak | GCA_008782695.1 | scaffold2220 (minus)SJXU01029907.1 | >ubap2l region: 605627-568842 | >hax1 XM_065903593.1 | >aqp10 region: 549076-546948 | >atp8b2 region: 540607-520794 | >il6r region: 463664-433709 |
| 22 | Reeves' muntjac | Muntiacus reevesi | GCA_963930625.1 | Chr. 1 (minus) | >ubap2l XM_065903440.1 | | >aqp10 XM_065945803.1 | >atp8b2 XM_065903394.1 | >il6r XM_065903394.1 |
| 23 | Reindeer | Rangifer tarandus | GCA_949782905.1 | scaffold2245 (minus)JAJJMQ010013826.1 | >ubap2l region: 286712-251599 | >hax1 region: 15477405-15475646 | >aqp10 region: 228831-226735 | >atp8b2 region: 221675-203617 | >il6r region: 142449-107529 |
| 24 | Mule deer | Odocoileus hemionus | GCA_020976825.1 | Chr. 1 (minus)JAJLRB010000001.1 | >ubap2l region: 15516276-15481194 | >hax1 XM_020887373.2 | >aqp10 region: 15460051-15458011 | >atp8b2 region: 15453397-15437021 | >il6r region: 15379738-15350188 |
| 25 | White-tailed deer | Odocoileus virginianus | GCA_023699985.4 | Chr. 5 (plus) | >ubap2l XM_020887354.2 | | >aqp10 XM_070467125.1 | >atp8b2 XM_020887377.2 | >il6r XM_020887383.2 |
| 26 | Cattle | Bos taurus | GCA_002263795.4 | Chr. 3 (minus) | >ubap2l NM_001045953.2 | >hax1 XM_005898646.2 | >aqp10 XM_024989821.1 | >atp8b2 NM_001191221.2 | >il6r NM_001110785.3 |
| 27 | Wild yak | Bos mutus | GCA_027580195.2 | Chr. 3 (plus) | >ubap2l XM_070367219.1 | >hax1 XM_006044198.3 | >aqp10 XM_005898757.2 | >atp8b2 XM_005898648.2 | >il6r XM_005898650.2 |
| 28 | Water buffalo | Bubalus bubalis | GCA_019923935.1 | Chr. 6 (minus) | >ubap2l XM_025287759.2 | >hax1 XM_010838351.1 | >aqp10 XM_006044200.3 | >atp8b2 XM_044944309.1 | >il6r XM_006044204.3 |
| 29 | American bison | Bison bison | GCA_000754665.1 | NW_011494665.1 (minus) | >ubap2l XM_010838353.1 | >hax1 XM_042254637.1 | >aqp10 XM_010838441.1 | >atp8b2 XM_010838347.1 | >il6r XM_010838440.1 |
| 30 | Sheep | Ovis aries | GCA_016772045.2 | Chr. 1 (plus) | >ubap2l XM_012182307.5 | >hax1 XM_069575306.1 | >aqp10 XM_004003657.5 | >atp8b2 XM_012182307.5 | >il6r XM_027975656.3 |
| 31 | Bighorn sheep | Ovis canadensis | GCA_042477335.2 | Chr. 1 (plus) | >ubap2l XM_069574927.1 | >hax1 XM_005677436.3 | >aqp10 XM_069575396.1 | >atp8b2 XM_069575409.1 | >il6r XM_069575409.1 |
| 32 | Goat | Capra hircus | GCA_001704415.2 | Chr. 3 (plus) | >ubap2l XM_013976959.2 | >hax1 region: 17115507-17113731 | >aqp10 XM_005677552.2 | >atp8b2 XM_018046190.1 | >il6r XM_018046192.1 |
| 33 | Muskox | Ovibos moschatus | GCA_041156055.1 | scaffold260(minus) JACAUE020000305.1 | >ubap2l region: 17155310-17117918 | >hax1 region: 104761968-104763740 | >aqp10 region: 17097492-17095373 | >atp8b2 region: 17090837-17074451 | >il6r region: 17011454-16979588 |
| 34 | Chiru | Pantholops hodgsonii | GCA_040182635.1 | Chr. 5 (plus)JBBYXG010000005.1 | >ubap2l region: 104722099-104759562 | >hax1 XM_040230482.1 | >aqp10 region: 104780189-104782310 | >atp8b2 region: 104786825-104802950 | >il6r region: 104866764-104898436 |
| 35 | Scimitar-horned oryx | Oryx dammah | GCA_014754425.2 | NW_024070212.1 (minus) | >ubap2l XM_040230451.1 | | >aqp10 XM_040229914.1 | >atp8b2 XM_040230442.1 | >il6r XM_040230441.1 |
| 36 | Chinese forest musk deer | Moschus berezovskii | GCA_022376915.1 | NW_026574040.1 (plus) | >ubap2l XM_055404107.1 | >hax1 region: 7630030-7628256 | >aqp10 XM_055403765.1 | >atp8b2 XM_055402951.1 | >il6r XM_055402067.1 |
| 37 | Siberian musk deer | Moschus moschiferus | GCA_004024705.2 | BS20 Scw3uUT_1114165;HRSCAF=1217686_2 (minus) PVHU021072402.1 | >ubap2l region:7671024-7632399 | | >aqp10 region: 7611680-7609136 | >atp8b2 region: 7604690-7588313 | >il6r region: 7530511-7499950 |
| 38 | Hippopotamus | Hippopotamus amphibius | GCA_030028045.1 | Chr. 1 (plus) | >ubap2l XM_057725330.1 | >hax1 region: 108866395-108868191 | >aqp10 XM_057716216.1 | >atp8b2 XM_057725368.1 | >il6r XM_057725373.1 |
| 39 | Pygmy hippopotamus | Choeropsis liberiensis | GCA_023065765.1 | Chr. 1 (plus)JAKVTR010000001.1 | >ubap2l region: 108829040-108863922 | | >aqp10 region: 108882991-108885449 | >atp8b2 region: 108890086-108907330 | >il6r region: 108967382-108993819 |
| 40 | Minke whale | Balaenoptera acutorostrata | GCA_949987535.1 | Chr. 1 (plus) | >ubap2l XM_057528496.1 | >hax1 XM_036823641.1 | >aqp10 XM_007198353.1 | >atp8b2 XM_057555548.1 | >il6r XM_007178486.3 |
| 41 | Blue whale | Balaenoptera musculus | GCA_009873245.3 | Chr. 1 (plus) | >ubap2l XM_036857869.1 | >hax1 XM_068540897.1 | >aqp10 XM_036867377.1 | >atp8b2 XM_036873778.1 | >il6r XM_036831078.1 |
| 42 | Grey whale | Eschrichtius robustus | GCA_028021215.1 | Chr. 3 (plus) | >ubap2l XM_068540880.1 | | >aqp10 XM_068537587.1 | >atp8b2 XM_068540903.1 | >il6r XM_068540905.1 |
| 43 | Sperm whale | Physeter macrocephalus | GCA_002837175.5 | NW_021146240.1 (minus) | | >hax1 XM_060007865.1 | >aqp10 XM_007130325.1 | | |
| 44 | Saddleback dolphin | Delphinus delphis | GCA_949987515.2 | Chr. 1 (plus) | >ubap2l XM_060030509.1 | >hax1 XM_004330551.4 | >aqp10 XM_060007887.1 | >atp8b2 XM_060030649.1 | >il6r XM_060030656.1 |

| | | | | | | | | | |
|---|---|---|---|---|---|---|---|---|---|
| 45 | Bottlenosed dolphin | Tursiops truncatus | GCA_011762595.2 | Chr. 1 (minus) | >ubap2l XM_033858292.2 | >hax1 XM_067727943.1 | >aqp10 XM_004330549.4 | >atp8b2 XM_033858129.2 | >il6r XM_019919452.3 |
| 46 | False Killer Whale | Pseudorca crassidens | GCA_039906515.1 | Chr. 2 (plus) | >ubap2l XM_067727917.1 | >hax1 XM_060137774.1 | >aqp10 XM_067727951.1 | >atp8b2 XM_067727911.1 | >il6r XM_067727965.1 |
| 47 | White-beaked dolphin | Lagenorhynchus albirostris | GCA_949774975.1 | Chr. 2 (minus) | >ubap2l XM_060137472.1 | >hax1 XM_004284661.3 | >aqp10 XM_060137809.1 | >atp8b2 XM_060137408.1 | >il6r XM_060137686.1 |
| 48 | Killer whale | Orcinus orca | GCA_937001465.1 | Chr. 1 (minus) | >ubap2l XM_033414575.2 | | >aqp10 XM_004284659.3 | >atp8b2 XM_049712250.1 | >il6r XM_049697651.1 |
| 49 | Harbour Porpoise | Phocoena phocoena | GCA_963924675.1 | Chr. 1 (plus) | >ubap2l XM_065880315.1 | >hax1 XM_022559773.2 | >aqp10 XM_065895752.1 | >atp8b2 XM_065897102.1 | >il6r XM_065895832.1 |
| 50 | Beluga whale | Delphinapterus leucas | GCA_002288925.3 | XM_065895832.1 (plus) | >ubap2l XM_030759555.1 | >hax1 XM_007457996.1 | >aqp10 XM_022559619.2 | >atp8b2 XM_022559374.1 | >il6r XM_022559475.2 |
| 51 | Yangtze River dolphin | Lipotes vexillifer | GCA_000442215.2 | NW_006785665.1 (minus) | >ubap2l XM_007458004.1 | | >aqp10 XM_007457995.1 | >atp8b2 XM_007457992.1 | >il6r XM_007457991.1 |
| 52 | Horse | Equus caballus | GCA_041296265.1 | Chr. 5 (minus) | >ubap2l XM_070268133.1 | | >aqp10 XM_023641093.2 | >atp8b2 XM_023641090.2 | >il6r XM_005610076.3 |
