## Supplementary material for "Evolutionary history of aquaporin-10 pseudogenization in Cetartiodactyla": Fig. S2

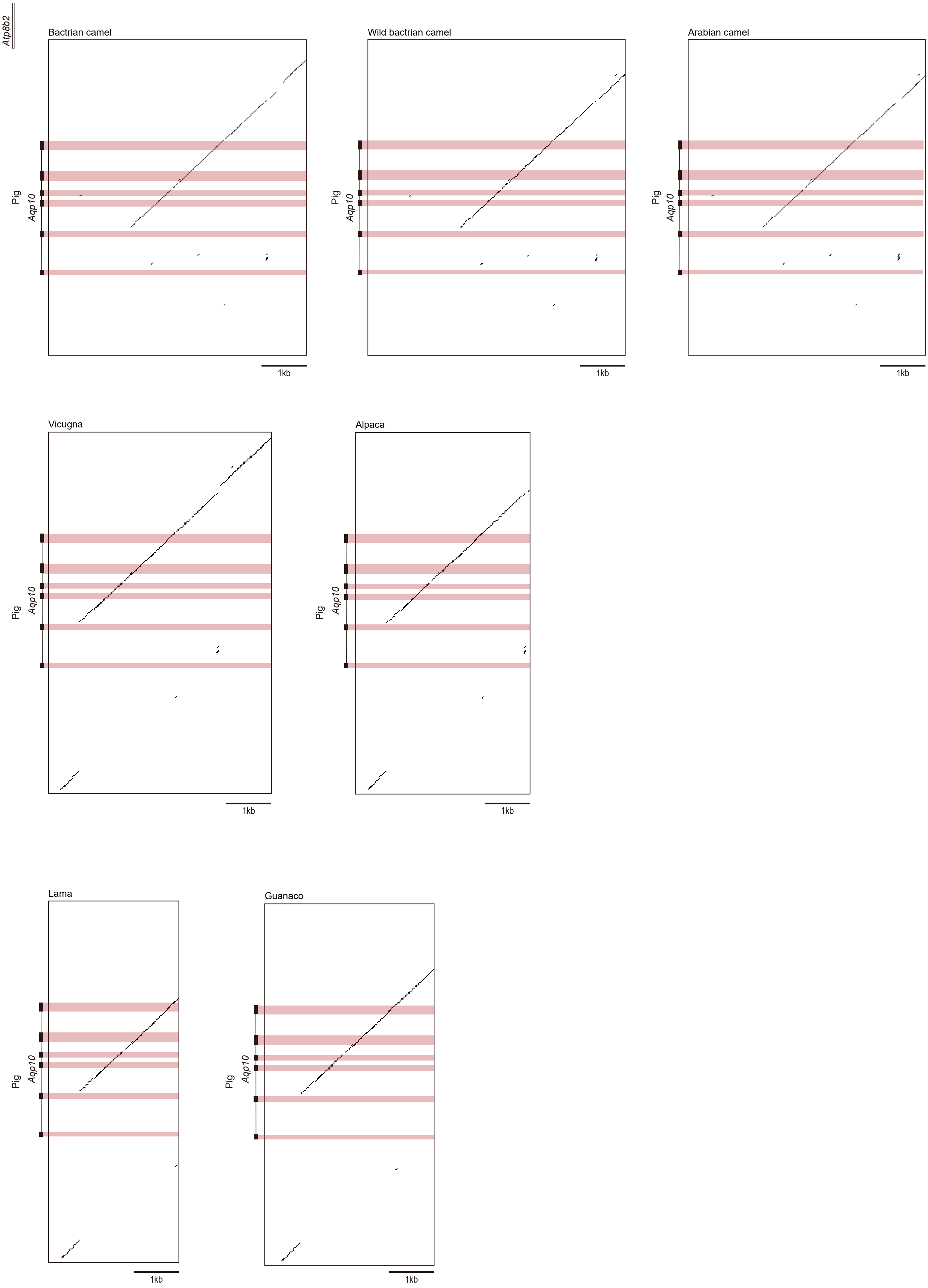

Atp8b2
Bactrian camel
Wild bactrian camel
Arabian camel
Vicugna
Alpaca
Lama
Guanaco
Pig
Aqp10
1kb

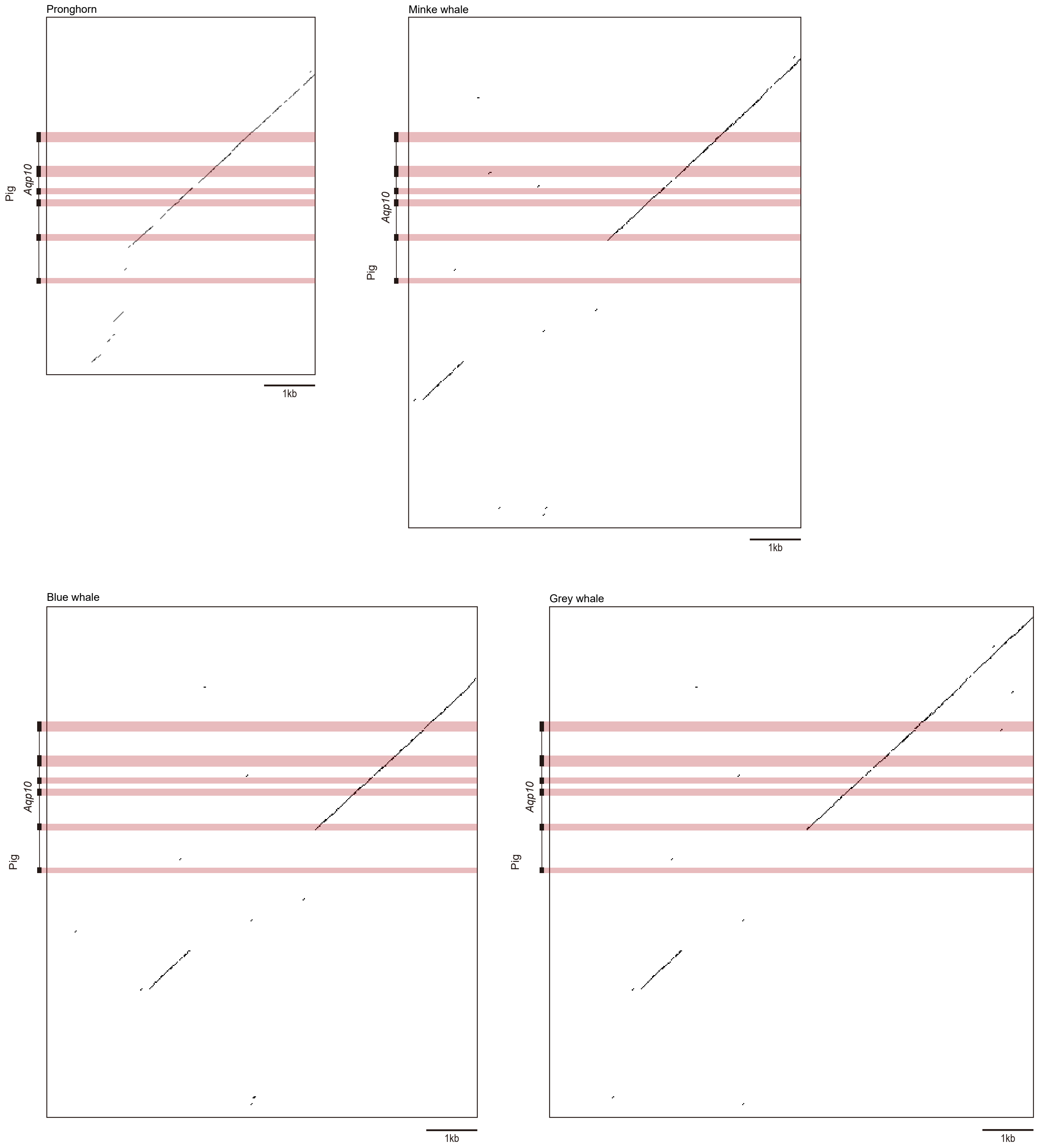


**Fig. S2. Dot plot analyses of *Aqp10* pseudogenes in Artiodactyla.**
The *Aqp10* gene and its flanking regions in comparison with the corresponding genome regions of pig various Artiodactyla species are shown. Homologous regions were plotted with the EMBOSS dotmatcher program (https://www.ebi.ac.uk/jdispatcher/emboss) with a window size of 20 and threshold score of 70. The genomic regions containing the analyzed Aqp10p loci are listed in Table S2.
