## Supplementary material for "Evolutionary history of aquaporin-10 pseudogenization in Cetartiodactyla": Table S1

**Table S1.** Manually predicted nucleotide and amino acid sequences of Cetartiodactyla AQP10

| No. | AQP10 sequences | Size |
|---|---|---|
| 3 | **North Sulawesi babirusa (*Babyrousa celebensis*)** | |
| | atggcctacactctgtccctggccaaactcaaggaacggctccggatcggcaacctcctggtccggcagtgcctggcagagtttctaggtgtgtttgtgctcatgctcttcacccaggagtctgtggcccaggcagtaaccagtggagaaactaaaggcaacttctttaccatgtttctggctggcgctctgggcgttatactagccatctacattggtggtaatgtctcagggggccacctgaatccagccttctccctggccatgtgtctcttgggacgcttcccctgggccaagtttcccatttactccttggtgcagttggtgtctgctttctgtgcctctgcagccacctatgcactctattacgatgccctacagaactatacaggtgggaacctgacagtgactggtcccaaggagacagcctccatctttgccacctatcctgccccttatctatccttgagcaatggcatccttgatcaggttctgggcacctggttgctgatggtggggatctcagctatcctggacacacggaacaagggagtgcctgcaggtctggagcctgtggtagtgggattgctgatcctggccattgggctatccatgggtgtcaactgtgggtacccgatgaaccctgcccgggacctgggcccacggctcttcacctatgtggccggctggggtcctgaagtcttcagcgctggtaatggttggtggtgggtgcctgtggtgtgtcctctggtgggggctgctcttggcacaaccacatatcagctgctggtggctctgcaccatactgaggactcagagacagctcaggaagtaaactgtgcccaacctgaagtctcagcctga | 864 nt |
| | MAYTLSLAKLKERLRIGNLLVRQCLAEFLGVFVLMLFTQESVAQAVTSGETKGNFFTMFLAGALGVILAIYIGGNVSGGHLNPAFSLAMCLLGRFPWAKFPIYSLVQLVSAFCASAATYALYYDALQNYTGGNLTVTGPKETASIFATYPAPYLSLSNGILDQVLGTWLLMVGISAILDTRNKGVPAGLEPVVVGLLILAIGLSMGVNCGYPMNPARDLGPRLFTYVAGWGPEVFSAGNGWWWVPVVCPLVGAALGTTTYQLLVALHHTEDSETAQEVNCAQPEVSA | 287 aa |
| 4 | **Chacoan peccary (*Catagonus wagneri*)** | |
| | atggcctacactctgtccctggccaaactcaaggaccggctccggatccacagcctcctggtccgagagtgcctggcggagttcctaggtgtgtttgtgctcatgctcttcacccagggatctgtggcccaggctgtcaccagtggagaaactaaaggcaacttcttcaccatgtttctggctggcgctctgggcgttatcgtagccatctacgtgggagggaatgtctcagggggccacctgaatccagccttctccctggccatgtgtctcctgggacgcctcccctgggccaagtttcccatttactccttggtgcagttggtgtctgccttctgtgcctctggagccacctatgcactctattatgatgccctacagaactatacaggtgggaacctgacagtgactggtcccaaggagacagcctccatctttgccacctatcctgtcccttatctatctttgagcaatggcatcctggatcaggttcttggcacctggatgctgattgtggggatctcggctatcctggacacacggaacaagggagtgcctgcgggtctggagcctgtggtagtggggttgctgatcctggccatcgggctatccatgggtgtcaactgtgggtacccaatgaaccctgcccgggacctgggcccacggctcttcacctatgtggccggctggggccctgaagtcttcagcgctggaaatggctggtggtgggtgcctgtggtgggccctctggtgggggccactcttggcacagccacatatcagctgctggtggctctgcattatcctgaggactcagagccagctcaggaactaaagtgtgcccaacctgaagtgtcaggggcaacacttcagcggacaactcggatgcggacgagtaagctgtga | 906 nt |
| | MAYTLSLAKLKDRLRIHSLLVRECLAEFLGVFVLMLFTQGSVAQAVTSGETKGNFFTMFLAGALGVIVAIYVGGNVSGGHLNPAFSLAMCLLGRLPWAKFPIYSLVQLVSAFCASGATYALYYDALQNYTGGNLTVTGPKETASIFATYPVPYLSLSNGILDQVLGTWMLIVGISAILDTRNKGVPAGLEPVVVGLLILAIGLSMGVNCGYPMNPARDLGPRLFTYVAGWGPEVFSAGNGWWWVPVVGPLVGATLGTATYQLLVALHYPEDSEPAQELKCAQPEVSGATLQRTTRMRTSKL | 301 aa |
| 12 | **Java mouse-deer (*Tragulus javanicus*)** | |
| | atggtctacattcagtccctggcggaggtcaaggactggctccggctccgcagcctcctggcccggcagtgcctggcggagttcctgggtgtgtttgtgctcatgctcctcatacaggggcttcggcccaggctgtcaccagtggaggaaccaaaggcaacttcttcaccatgtttctggccggctctctgggtgtggtggtggccatctatgtgagtggtaatgtctcaggggcccacttgaatccagccttctccctggccatgtgcctcctgggacgcctcccctgggccacgtttcccgtttactccttggtgcagttgctgtctgctttctgtgcctctggagtcacctttgctctctattatgatgccctacagaactatacaggtgggaacctgaccgtgacaggtcccaaggagacagcctccatctttgccacctaccctgcccccctatctgtccctgagcaacggcttcctggatcaggttctgggcacctggatgctgatcgtggggatcttggccatcctggacacacggaacaagggagtgcctgcaggtctggagcctgtggtagtggggttgctgatcctggccatcgcactctccatgggtgccaactgtgggctcccaatcaaccccgcccgcgacctcggcccacggcttttcacctacgtggctggctggggccccgaagtattcagcgctggccacggctggtggtgggtgcctgtggtgggtcctctagtgggagccacacttggcacagccacataccagctgctggtggccctgcaccatcctgaggactcagagccagctcaggatctaaagggtgcccaaccggaagcctcagattcgggaagcgctgccgcgactcccgtgcaggagagtaagctgtaa | 906 nt |
| | MVYIQSLAEVKDWLRLRSLLARQCLAEFLGVFVLMLLIQGASAQAVTSGGTKGNFFTMFLAGSLGVVVAIYVSGNVSGAHLNPAFSLAMCLLGRLPWATFPVYSLVQLLSAFCASGVTFALYYDALQNYTGGNLTVTGPKETASIFATYPAPYLSLSNGFLDQVLGTWMLIVGILAILDTRNKGVPAGLEPVVVGLLILAIALSMGANCGLPINPARDLGPRLFTYVAGWGPEVFSAGHGWWWVPVVGPLVGATLGTATYQLLVALHHPEDSEPAQDLKGAQPEASDSGSAAATPVQESKL | 301 aa |
| 13 | **Lesser mouse-deer (*Tragulus kanchil*)** | |
| | atggtctacattcagtccctggcggaggtcaaggactggctccggctccgcagcctcctggcccggcagtgcctggcggagttcctgggtgtgtttgtgctcatgctcctcatacaggggcttcggcccaggctgtcaccagtggaggaaccaaaggcaacttcttcaccatgtttctggccggctctctgggtgtggtggtggccatctatgtgagtggtaatgtctcaggggcccacttgaatccagccttctccctggccatgtgcctcctgggacgcctcccctgggccacgtttcccatttactccttggtgcagttgctgtctgctttctgtgcctctggagtcacctttgctctctattatgatgccctacagaactatacaggtgggaacctgaccgtgacaggtcccaaggagacagcctccatctttgccacctaccctgcccccctatctgtccctgagcaacggcttcctggatcaggttctgggcacctggatgctgatcgtggggatcttggccatcctggacacacggaacaagggagtgcctgcaggtctggagcctgtggtagtggggttgctgatcctggccatcgcactctccatgggtgccaactgtgggctcccaatcaaccccgcccgcgacctcggcccacggcttttcacctacgtggctggctggggccccgaagtcttcagcgctggccacggctggtggtgggtgcctgtggtgggtcctctagtgggagccacacttggcacagccacataccagctgctggtggccctgcaccatcctgaggactcagagccagctcaggatctaaagggtgcccaaccggaagcctcagattcgggaagcgctgccgcgactcccgtgcaggagagtaagctgtaa | 906 nt |
| | MVYIQSLAEVKDWLRLRSLLARQCLAEFLGVFVLMLLIQGASAQAVTSGGTKGNFFTMFLAGSLGVVVAIYVSGNVSGAHLNPAFSLAMCLLGRLPWATFPIYSLVQLLSAFCASGVTFALYYDALQNYTGGNLTVTGPKETASIFATYPAPYLSLSNGFLDQVLGTWMLIVGILAILDTRNKGVPAGLEPVVVGLLILAIALSMGANCGLPINPARDLGPRLFTYVAGWGPEVFSAGHGWWWVPVVGPLVGATLGTATYQLLVALHHPEDSEPAQDLKGAQPEASDSGSAAATPVQESKL | 301 aa |
| 16 | **Rothschild's giraffe (*Giraffa camelopardalis rothschildi*)** | |
| | atggactccattcagtctctggttgaagtcaaggaccggctccagatctgcagcctcctggcccagcagtgcctggcggagtttctggctgtgtttgtgctcatactcctcatacaagggtcttcggcccaggctgtcaccagtggaggaaccaaaggaaacttctccaccatgtttctggctggctctctgagcgttatgctagccatctatgtgactggtaatgtctcaggggcccacctgaatccagccttctccctggccatgtgtctcctgggatgcctcccccgggccaagtttctcatttactccttggtgcagttgctgtctgccttctgtgcttctggagtcacctatgctctgtattatgatgccctacagaactacacaggtgggaacctgacagtgactggtcccaaggagacagcttccatctttgccacctaccctgcccccctatctgtccctgagcaatggcttcctggatgaggttctgggcacctggatgctgatggtggggatcctggccatcctggacacacggaacaagggagtgcctgcgggtctggagcctgtggtagtggggttactgatcctggccatcacactatccgtgggtgccaactgtgggctcctaatcaaccctgcccaggacctcggcccagggcttttcacctacgtagctggctgtggccctgaagtcttcaatgctggctatggttggtggtgggtccctgtggtgggccctctagtgggggccatgctaggcccagccacataccagctgctggtggcacagcaccatcctgaggactcagagccatctcaggatctagagtgtgcccaacaggaagcctcagactcgggaagtccggccgcgactcagctgcaggagagtaagctgtga | 906 nt |
| | MDSIQSLVEVKDRLQICSLLAQQCLAEFLAVFVLILLIQGSSAQAVTSGGTKGNFSTMFLAGSLSVMLAIYVTGNVSGAHLNPAFSLAMCLLGCLPRAKFLIYSLVQLLSAFCASGVTYALYYDALQNYTGGNLTVTGPKETASIFATYPAPYLSLSNGFLDEVLGTWMLMVGILAILDTRNKGVPAGLEPVVVGLLILAITLSVGANCGLLINPAQDLGPGLFTYVAGCGPEVFNAGYGWWWVPVVGPLVGAMLGPATYQLLVAQHHPEDSEPSQDLECAQQEASDSGSPAATQLQESKL | 301 aa |
| 37 | **Siberian musk deer (*Moschus moschiferus*)** | |
| | atggcatggctttgctccatagccctcaaggaccggctccgtatctgcaggctcctggcccggcagtgcctggcagagtttctggctgtgtttgtgctcatggtagggtctttggcccaggctgtcaccagtgtaggaaccaaaggaaacttctccactgtgtttctggctggctctctgggcgttatgctagtcatctatgtgagtggtgatgtctcaggggcccacctgaatccagccttctccctgcccgtttgcctctggggacgcctcccctgggccacgtttctcatttactccttggtgcagttgctgtgtgcttcctgtgcctctggagtcacctatgctctctattacgatgccctacagaactacacaggtgggagcctgacagtgactggtcccaaggagatggcttccagctttgccacctaccctgcccccctatctgtctctgagcaatggcttcctggatcaggttctgggcacctggatgctaatggtggggatcctggctattctgaacacacggaacaagggagtgcctgcaggtctggagcctgtggtgttggggttactgatcctggccatcacgctatccatgggtgccaactgtgggctcccaatcaaccctgcccaggaccttggcccacggcttttcacctgcatagctggctggggccctgaagtcttcagtgctggcaatagttggtggtgggttcctgtggtgggccctctagtgggggccacgcttggcacagccacataccagctgctggtggcacagcaccatcctgaagactcagagccatgtcaggatctagagtgtgcccaacaggaagcctcagactcgggaagtccggctgcaactcagctacaggagagtaagctgtga | 894 nt |
| | MAWLCSIALKDRLRICRLLARQCLAEFLAVFVLMVGSLAQAVTSVGTKGNFSTVFLAGSLGVMLVIYVSGDVSGAHLNPAFSLPVCLWGRLPWATFLIYSLVQLLCASCASGVTYALYYDALQNYTGGSLTVTGPKEMASSFATYPAPYLSLSNGFLDQVLGTWMLMVGILAILNTRNKGVPAGLEPVVLGLLILAITLSMGANCGLPINPAQDLGPRLFTCIAGWGPEVFSAGNSWWWVPVVGPLVGATLGTATYQLLVAQHHPEDSEPCQDLECAQQEASDSGSPAATQLQESKL | 297 aa |
