## Supplementary material for "Evolutionary history of aquaporin-10 pseudogenization in Cetartiodactyla": Table S2

**Table S2.** *Aqp10* regions used for dot plot analysis.

| No. | Common name | Species | Accession no. | Direction | Region |
|---|---|---|---|---|---|
| 1 | pig | *Sus scrofa* | NC_010446.5 | minus | 95448000..95458000 |
| 5 | Arabian camel | *Camelus dromedarius* | NC_087458.1 | plus | 6242000..6247000 |
| 6 | Wild bactrian camel | *Camelus ferus* | NC_045716.1 | minus | 22661000..22670000 |
| 7 | Bactrian camel | *Camelus bactrianus* | NW_011513412.1 | minus | 2092000..2098000 |
| 8 | Vicugna | *Vicugna vicugna* | PNXW01026126.1 | minus | 88000..94000 |
| 9 | Guanaco | *Lama guanicoe cacsilensis* | PNXV01013028.1 | minus | 276000..281000 |
| 10 | Lama | *Lama glama chaku* | PNXU01039556.1 | plus | 5000..11000 |
| 11 | Alpaca | *Vicugna pacos* | NW_021964201.1 | plus | 6370000..6377000 |
| 14 | Pronghorn | *Antilocapra americana* | SRVM01010816.1 | minus | 247000..252000 |
| 40 | Minke whale | *Balaenoptera acutorostrata* | CATLKR010006866.1 | plus | 254000..263000 |
| 41 | Blue whale | *Balaenoptera musculus* | VNFD02003604.1 | plus | 87000..95000 |
| 42 | Grey whale | *Eschrichtius robustus* | NC_090826.1 | plus | 112363000..112372000 |
