## Supplementary material for "Evolutionary history of aquaporin-10 pseudogenization in Cetartiodactyla": Fig. S3

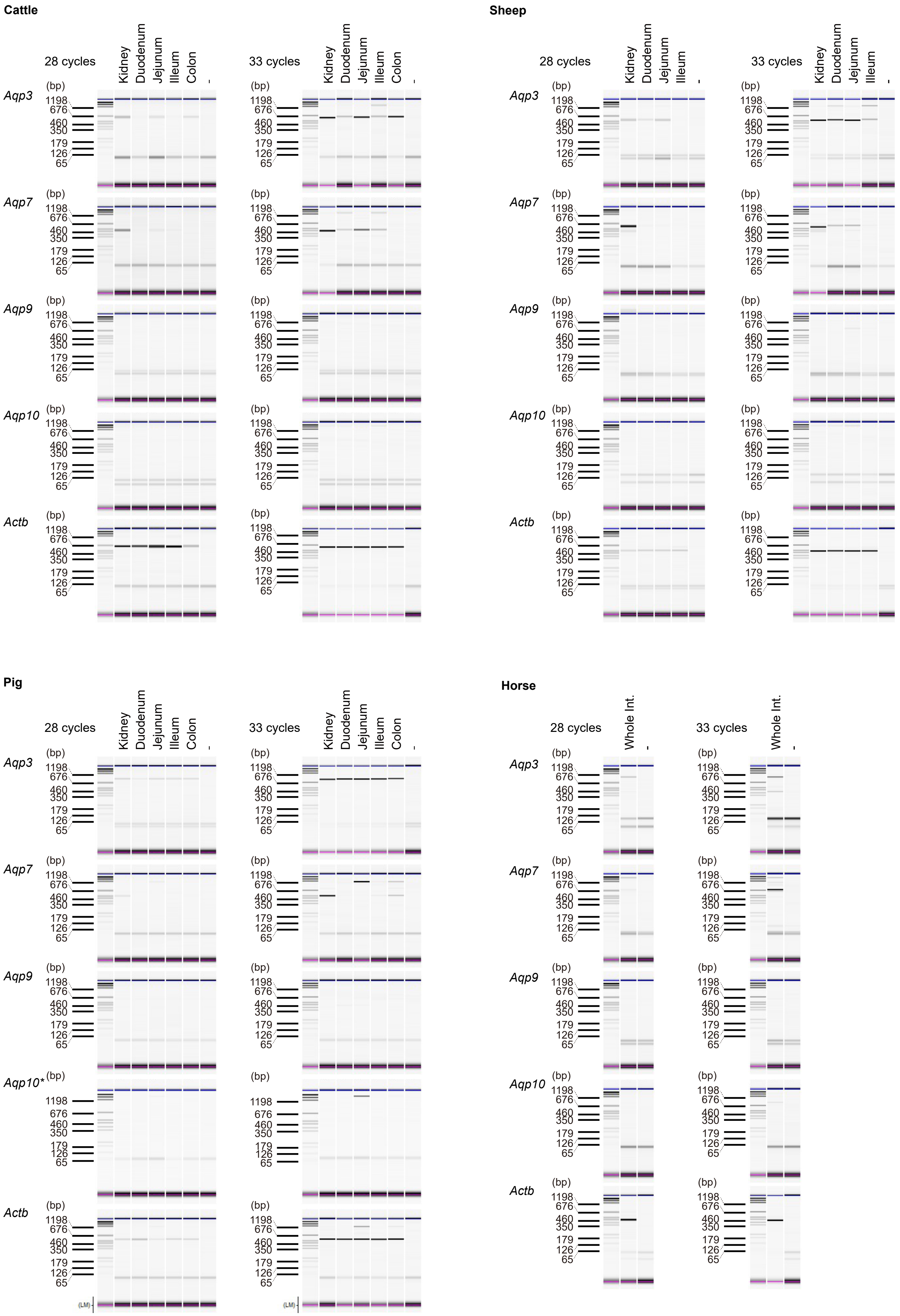


**Fig. S3. Whole images of a Microchip Electrophoresis system.**
Expression profiles of *Aqp10* and the other aquaporin genes in cattle, sheep, pig, and horse tissues were determined using semiquantitative RT-PCR. Pseudo-gel images of the PCR products were generated using a microchip electrophoresis system for DNA/RNA Analysis MCE-202 MultiNA. *: Annealing temperature of 56°C
